# Regrowth-delay Body as a Bacterial Subcellular Structure marking multidrug tolerant Persisters

**DOI:** 10.1101/448092

**Authors:** Jiayu Yu, Yang Liu, Huijia Yin, Zengyi Chang

**Affiliations:** The State Key Laboratory of Protein and Plant Gene Research, School of Life Sciences, Peking University, Beijing 100871, P.R. China; Center for Protein Science, Peking University, Beijing 100871, P.R. China

## Abstract

Bacteria have long been recognized to be capable of entering a phenotypically non-growing persister state, in which the cells exhibit an extended regrowth lag and a multidrug tolerance, thus posing a great challenge in treating infectious diseases. Owing to their non-inheritability, low abundance of existence, lack of metabolic activities, and high heterogeneity, properties of persisters remain poorly understood. Here, we report our accidental discovery of a hitherto unreported subcellular structure that we term the regrowth-delay body, which is formed only in non-growing bacterial cells and sequesters multiple key proteins. As of now, this structure, that dissolves when the cell resumes growth, is the most distinguishable subcellular structure marking persisters. Our studies also indicate that persisters exhibit different depth of persistence, as determined by the status of their regrowth-delay bodies. Our findings imply that suppressing the formation and/or promoting the dissolution of regrowth-delay bodies could be viable strategies for eradicating persisters.

## INTRODUCTION

It has been well documented that, in a genetically homogeneous population of bacterial cells, a subset are able to enter a phenotypically dormant, non-growing (or, more precisely, of low metabolic activity) state. This state has been variably named as sporulation, latency, regrowth lag, persisters, or the viable but nonculturable, in laboratory, clinical, or environmental microbiology (Burke et al., 1925; Chesney, 1916; Kaprelyants et al., 1993; Lewis, 2007, 2010; Monod, 1949; Roszak and Colwell, 1987). Although this state of bacterial cells has been recognized for more than 100 years, much remain unknown on its properties, such as how the bacterial cells enter, maintain and exit such a unique state, that is best known for its non-inheritable multidrug tolerance (Balaban et al., 2013; Kaldalu et al., 2016; Kell et al., 2015; Lewis, 2007; Pinto et al., 2015).

The regrowth lag phenomenon, initially recognized by Max Muller in 1895 (Chesney, 1916), was observed as soon as bacterial culturing became feasible (Coplans, 1910), but remains the most poorly understood stage of the bacterial growth cycle (Monod, 1949; Rolfe et al., 2012). In a related phenomenon, bacterial dormancy was defined as a state of certain bacterial cells that exhibits a long-lasting regrowth lag (Burke et al., 1925; Chesney, 1916). Later, the term persister was coined to denote an extremely small subpopulation of dormant, non-dividing bacterial cells that are not killed by concentrations of antibiotics sufficiently high to kill the actively dividing ones (Bigger, 1944). The persisters were presumed to be responsible for the post-treatment relapse of bacterial infections (Bigger, 1944; Fisher et al., 2017; Lewis, 2007, 2010; Mcdermott, 1958). It was emphasized that the persisters are not resistant to antibiotics, since they produce offspring that are as susceptible to antibiotics as their parent cells (Bigger, 1944). More recently, it was unveiled that the bacterial cells in the natural environment are commonly in a viable but nonculturable dormant state (Ayrapetyan et al., 2015; Xu et al., 1982), one that is highly similar to the persisters.

Although much effort has been made to understand the molecular mechanisms leading to the formation of persisters, and certain specific protein factors (like the Hip) or small molecules (like the pppGpp) have been claimed to be important for this process (Black et al., 1991, 1994; Moyed and Bertrand, 1983), not much is certain up to now (Balaban et al, 2013; Kaldalu et al, 2015; Korch et al., 2003; Chowdhury et al., 2016). The slow pace of learning about this state of bacterial cells is apparently attributed to the great technical difficulty of unequivocally identifying them, which are presumed to exist in extremely small numbers in a genetically uniform population, often with no significant morphological distinctions (Balaban et al., 2013; Kaldalu et al., 2016; Kell et al., 2015). Because of this, persisters have been hitherto commonly perceived only on the basis of their lack of growth and multidrug tolerance. In particular, persisters have been conventionally detected by indirectly measuring the number of colony-forming units (CFUs) after treating the cell samples with a high concentration of a certain antibiotic (Jiafeng et al., 2015; Orman and Brynildsen, 2015), or as cells that do not grow in the presence, but regrow after the removal, of antibiotics when monitored with a microfluidic device (Balaban et al., 2004).

We have been trying to explore proteins when they are present in living bacterial cells, as by performing protein photo-crosslinking analysis mediated by genetically introduced unnatural amino acids (Fu et al., 2013; Zhang et al., 2011). In one recent study, we examined the assembly patterns of the FtsZ protein, which plays an essential role by assembling into the Z-ring structure for each bacterial cell to divide into two via the cytokinesis process (Dai and Lutkenhaus, 1991; Erickson et al., 2010; Haeusser and Margolin, 2016), as well as for each mitochondrion (Beech et al., 2000) or chloroplast (TerBush et al., 2013) to divide into two. In particular, we revealed hitherto unreported lateral interactions between the FtsZ protofilaments that are essential for FtsZ to assemble into the dynamic Z-ring structure in living bacterial cells (Guan et al., 2018).

As an exciting byproduct of that study, we accidentally revealed the presence of a novel reversible subcellular structure that we named it as the regrowth-delay body. This structure is formed in non-growing late stationary-phase bacterial cells and sequesters multiple proteins essential for cell growth. Remarkably, the regrowth-delay bodies become dissolved when a bacterial cell exits the regrowth lag and resumes growth, meanwhile releasing the sequestered proteins for re-functioning. We also demonstrated that a higher degree of regrowth-delay body formation is correlated to a longer duration of regrowth lag as well as a higher level of antibiotic tolerance, not only in *E. coli* but also in two bacterial pathogens. Therefore, the regrowth-delay body not only acts as a unique and highly valuable biomarker for distinguishing the non-growing dormant persister cells from the actively growing non-persister cells, but also acts as a dynamic biological timer for bacterial cells to exit the regrowth lag. Our studies also indicate that each persister exhibits a particular depth of persistence, which seems to explain the long-observed heterogeneous nature of the persister subpopulation. Our findings should be proven greatly valuable not only for specifically identify and explore the persisters in any cell population, but also for designing viable strategies to eradicate the formidable multidrug-tolerant pathogenic persisters.

## RESULTS

### The cell division protein FtsZ no longer self-assembles but exists as an insoluble form in non-growing bacterial cells

In an attempt to unveil how FtsZ assembles into the dynamic Z-ring structure during the cytokinesis of bacterial cell division, we performed systematic protein photo-crosslinking analyses with FtsZ variants containing the genetically introduced photoactive unnatural amino acid pBpa (*p*-benzoyl-L-phenylalanine) (Chin et al., 2002) in living bacterial cells. This allowed us to uncover novel lateral interactions between the FtsZ protofilaments that were demonstrated to be essential for cell division (Guan et al., 2018).

During these studies, out of curiosity, we additionally examined the status of FtsZ in non-dividing/non-growing bacterial cells, as has never been addressed by people working with FtsZ. We revealed, as expected, that a FtsZ variant, though self-assembled into homo-oligomers in actively dividing cells (**Fig. S1A**, lanes 2 and 6), no longer does so (**Fig. S1A**, lanes 4 and 8) in the non-dividing/non-growing cells. Astonishingly, we observed that most of the free FtsZ monomers, together with almost all the photo-crosslinked products, were detected in the insoluble pellet fraction of lysates of the non-growing cells (**Fig. S1B**, lane 8). By contrast, all the photo-crosslinked FtsZ dimers and the free FtsZ monomers were principally detected in the soluble supernatant fractions of lysates of actively dividing cells (**Fig. S1B**, lane 3).

In light of this puzzling observation, we then examined the distribution pattern of the endogenous FtsZ (instead of the FtsZ variant we examined above) in *E. coli* cells. Likewise, we revealed that the endogenous FtsZ protein was largely detected in the soluble supernatant fraction of actively dividing cells (**Fig. 1A**, lane 2), but in the insoluble pellet fraction of the non-dividing/non-growing cells (lane 6). As comparison, we demonstrated that EF-Tu (one of the most abundant proteins in bacterial cells) and GroEL (a molecular chaperone binding to misfolded client proteins) were both largely detected in the supernatant fraction (**Fig. 1A**, lanes 2 and 5), with hardly any in the pellet fraction (lanes 3 and 6) of either actively dividing or non-dividing cells. Taken together, these results revealed for the first time that the FtsZ protein (as well as proteins interacting with it) exists as an insoluble form in non-dividing/non-growing bacterial cells.

**Figure 1.**
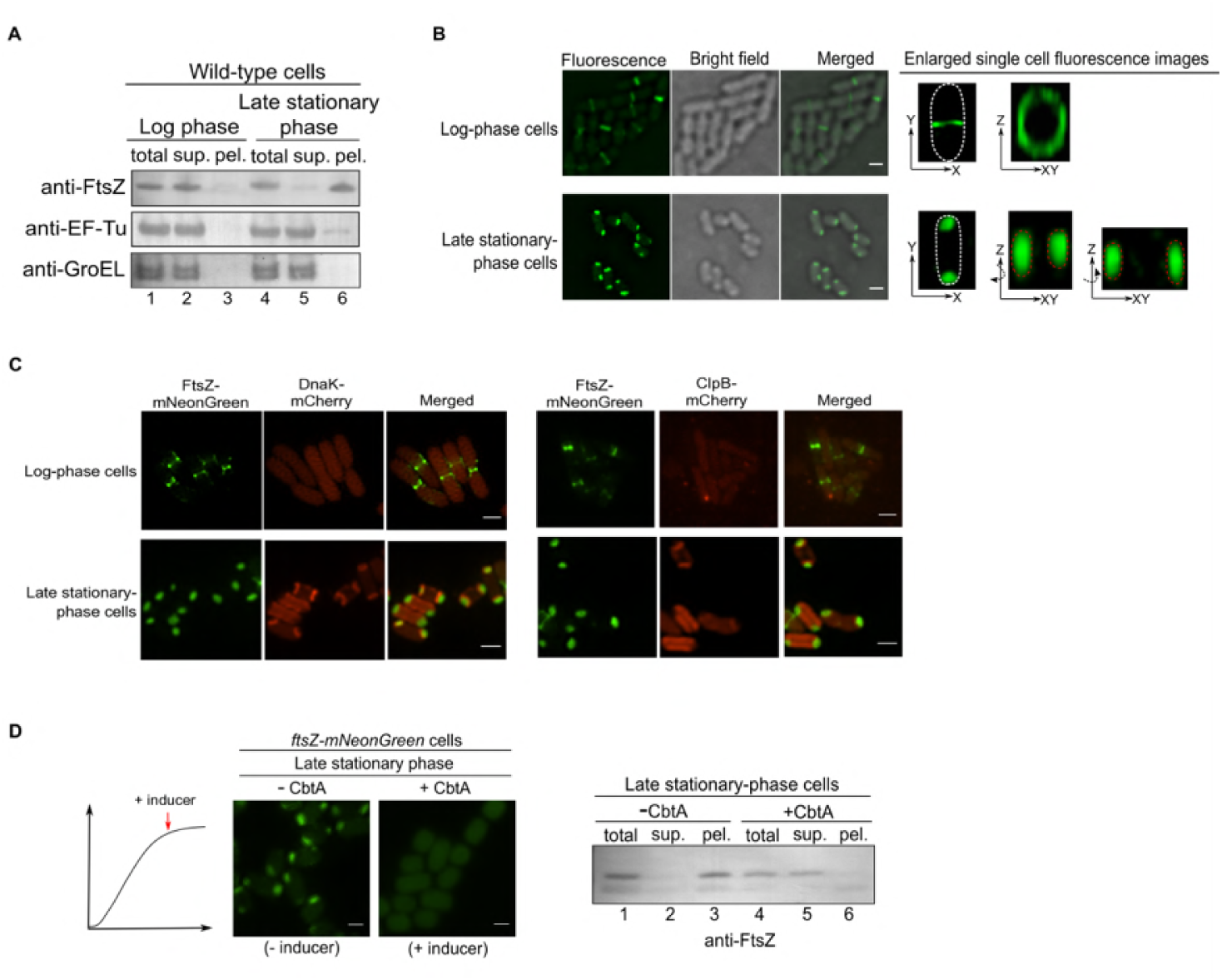
The cell division protein FtsZ in non-growing bacterial cells exists in a hitherto unreported cell-pole granule as a folded form. **(A)** Immunoblotting results for detecting endogenous FtsZ, EF-Tu, or GroEL in the total cell lysate (total), supernatant (sup.) and pellet (pel.) of the actively dividing logphase or the non-growing late stationary-phase wild-type *E. coll* cells, probed with the indicated antibodies. (See also **Fig. S1**) **(B)** Fluorescence and bright field microscopic images of the actively dividing log-phase (top) and the non-growing late stationary-phase (bottom) *E. coll* cells in which FtsZ-mNeonGreen was heterologously expressed. Scale bars, 1 μm. (See also *Figs. S2, S3*) **(C)** Fluorescence microscopic images of the actively dividing log-phase (top) and the non-growing late stationary-phase (bottom) *ftsZ-mNeonGreen-dnaK-mCherry* or *ftsZ-mNeonGreen-clpB-mCherry* cells. Scale bars, 1μm. **(D)** Fluorescence microscopic images of the non-growing late stationary-phase *ftsZ-mNeonGreen* cells in which the FtsZ inhibitor protein CbtA was expressed (left); the corresponding immunoblotting results for detecting FtsZ in the indicated cell lysate fractions, probed with anti-FtsZ antibodies (right). Scale bars, 1 μm. (See also **Fig. S4B-C**)

### The FtsZ protein exists in two cell-pole granules in each non-growing bacterial cell

We subsequently tried to monitor the status of FtsZ by performing live-cell imaging analysis. For this purpose, we started by heterologously expressing FtsZ-mNeonGreen, a form of FtsZ being fused to the green fluorescent protein mNeonGreen, in bacterial cells. Here, the fusion protein was expressed at a relatively low level, which was achieved via the leaky transcription of the Tet promoter (i.e., with no addition of the inducing agent), such that the fluorescent FtsZ fusion protein would be incorporated into, but not interrupt, the Z-ring structure that was largely formed via the assembly of endogenous wild type FtsZ. We first verified an effective incorporation of FtsZ-mNeonGreen into the Z-ring structure in actively dividing log-phase cells (**Fig. 1B**, top), like what was reported before (Ma et al., 1996). Remarkably, we then detected FtsZ-mNeonGreen as two cell pole-granules in each non-dividing cell (**Fig. 1B**, bottom). As a control, the unfused fluorescent mNeonGreen protein was shown to be evenly distributed in the cytoplasm of either actively-dividing or non-dividing bacterial cells (**Fig. S2**).

For further systematic live-cell imaging analysis, we subsequently constructed a bacterial strain whose genome was modified to express FtsZ-mNeonGreen (rather than from a plasmid), in parallel with the normally expressed endogenous FtsZ. In particular, we integrated the *ftsZ*-*mNeonGreen* gene into the genomic rhamnose operon (as illustrated in **Fig. S3A**) and demonstrated that the FtsZ-mNeonGreen protein would be produced only in the presence of rhamnose (the inducing sugar) in this *ftsZ-mNeonGreen* strain (**Fig. S3B**), hardly affecting the growth of the cells (**Fig. S3C**). We also verified the presence of FtsZ-mNeonGreen in the Z-ring structure in actively dividing log-phase but in the cell-pole granules in non-dividing late stationary-phase *ftsZ*-*mNeonGreen* cells (**Fig. S3D**).

Our live-cell imaging analysis employing this *ftsZ*-*mNeonGreen* strain revealed that the cell-pole granules seem to be closely associated with the inner membrane but not surrounded by it (**Fig. S4A**, middle panel), as verified by results (**Fig. S4B**) of staining with the membrane-specific dye FM4-64 (Fishov and Woldringh, 1999). These imaging results meanwhile demonstrated that the cell-pole granules occupy cytosolic locations that are hardly accessible to other cytosolic proteins (**Fig. S4A**, bottom panel), suggesting a compact nature. In line with this, we observed that these granules were maintained intact even after the cells were broken (**Fig. S4C**).

### The FtsZ protein in cell-pole granules are apparently folded

Aggregates of misfolded proteins have been reported to exist at the poles in *E. coli* cells, but only under heat shock conditions (Lindner et al., 2008; Winkler et al., 2010). Additionally, insoluble proteins, which were naturally assumed to be misfolded, have been reported to accumulate in stationary-phase *E. coli* cells (Kwiatkowska et al., 2008; Leszczynska et al., 2013; Maisonneuve et al., 2008). In view of these reports, we then attempted to clarify the folding status of FtsZ in the cell-pole granules, despite the fact that FtsZ was demonstrated to exist in a soluble form when heterologously over-expressed in bacterial cells (Mukherjee and Lutkenhaus, 1998).

Considering that the molecular chaperones DnaK and ClpB, as well as the protease ClpP were reported to be associated with protein aggregates formed under stress conditions (Winkler et al., 2010), we decided to analyze whether or not they are associated with the cell-pole granules. Our blotting analysis demonstrated that all these three quality control proteins were primarily detected in the supernatant (**Fig. S5A**, lane 2) with hardly any detected in the pellet (lane 3) of non-growing late stationary-phase cell lysates. In line with this, our live-cell imaging data showed that neither DnaK nor ClpB, each being expressed as a form fused to the red fluorescent protein mCherry (by manipulating their endogenous genes on the genomic DNA of the *ftsZ*-*mNeonGreen* strain), was detected in the cell-pole granules (**Fig. 1C**, bottom panels). The imaging data meanwhile revealed, interestingly, that both DnaK and ClpB, though being evenly dispersed in the cytosol of actively dividing log-phase cells, were concentrated near the two cell poles, at sites very close to but clearly separate from the FtsZ-containing cell-pole granules, but only in a small number of the non-growing late stationary-phase cells (**Fig. 1C**, bottom panels). These subcellular sites, which might represent ones where DnaK and ClpB (themselves being in soluble forms, as shown in **Fig. S5A**) were co-localized with certain form of protein aggregates, are worth further investigation in the future. Taken together, these results did not provide evidence to support the possibility that the cell-pole granules are typical aggregates formed by misfolded proteins.

As an attempt to further assess the folding status of FtsZ in the cell-pole granules, we examined whether inhibitor proteins that only bind to folded FtsZ could prevent FtsZ from entering the granules. For this purpose, we analyzed the CbtA and KilR proteins, each of which was known to bind to and to block monomeric FtsZ for assembling into the Z-ring structure in cells (Conter et al., 1996; Heller et al., 2017). Either CbtA or KilR was then expressed from a plasmid, under the control of an anhydrotetracycline-inducible promoter. We first verified their capacity to inhibit FtsZ from assembling into the Z-ring structure in actively dividing log-phase *ftsZ-mNeonGreen* cells (**Fig. S5B**).

We then showed that FtsZ was no longer able to enter the cell-pole granules when the CbtA expression was induced at the stationary phase (**Fig. 1D**, left panel). A similar effect was not observed when KilR was induced (**Fig. S5C**). In agreement with these findings, our immunoblotting analysis confirmed that FtsZ became undetectable in the pellet fraction but remained in the supernatant when CbtA was expressed (**Fig. 1D**, right panel).

Furthermore, the conclusion that FtsZ in the cell-pole granules is folded was also supported by our *in vivo* protein photo-crosslinking analysis. Specifically, the data (**Fig. S5D**) reveal that when the unnatural amino acid residue pBpa was placed at residue positions close to each other in space (e.g., positions 151, 166 and 174, or 31, 47, 51 and 54) according to the reported crystal structure (Löwe and Amos, 1998), similar patterns of photo-crosslinked products were generated. By contrast, when pBpa was placed at sites that were spatially distant (e.g., positions 61, 85, 299, and 340), different patterns of photo-crosslinked products were detected.

Collectively, these results strongly support the conclusion that FtsZ in the granules is folded, rather than misfolded.

### The cell-pole granules become dissolved in cells exiting their regrowth lag and resuming growth

We next sought to decipher the fate of the cell-pole granules when the bacterial cells resume their growth. For this purpose, we re-cultured the non-growing late stationary-phase *ftsZ-mNeonGreen* cells in fresh culture medium lacking the inducer rhamnose to avoid the production of new FtsZ-mNeonGreen protein. Remarkably, we observed an effective relocation of FtsZ-mNeonGreen from the cell-pole granules to the Z-ring structure that was formed in cells ending their regrowth lag and resuming the growth (**Fig. 2A**, exemplified by the cells circled with dashed pink or white lines). These newly assembled Z-ring structures seemed to be fully functional since they enabled the mother cells to split into two daughter cells (e.g., the cell circled with dashed pink lines at 80 min divided into two daughter cells at 120 min in **Fig. 2A**). Of equal importance, cells that remained in the regrowth lag state all retained their cell-pole granules (**Fig. 2A**, exemplified by the cells circled with red dashed lines).

**Figure 2.**
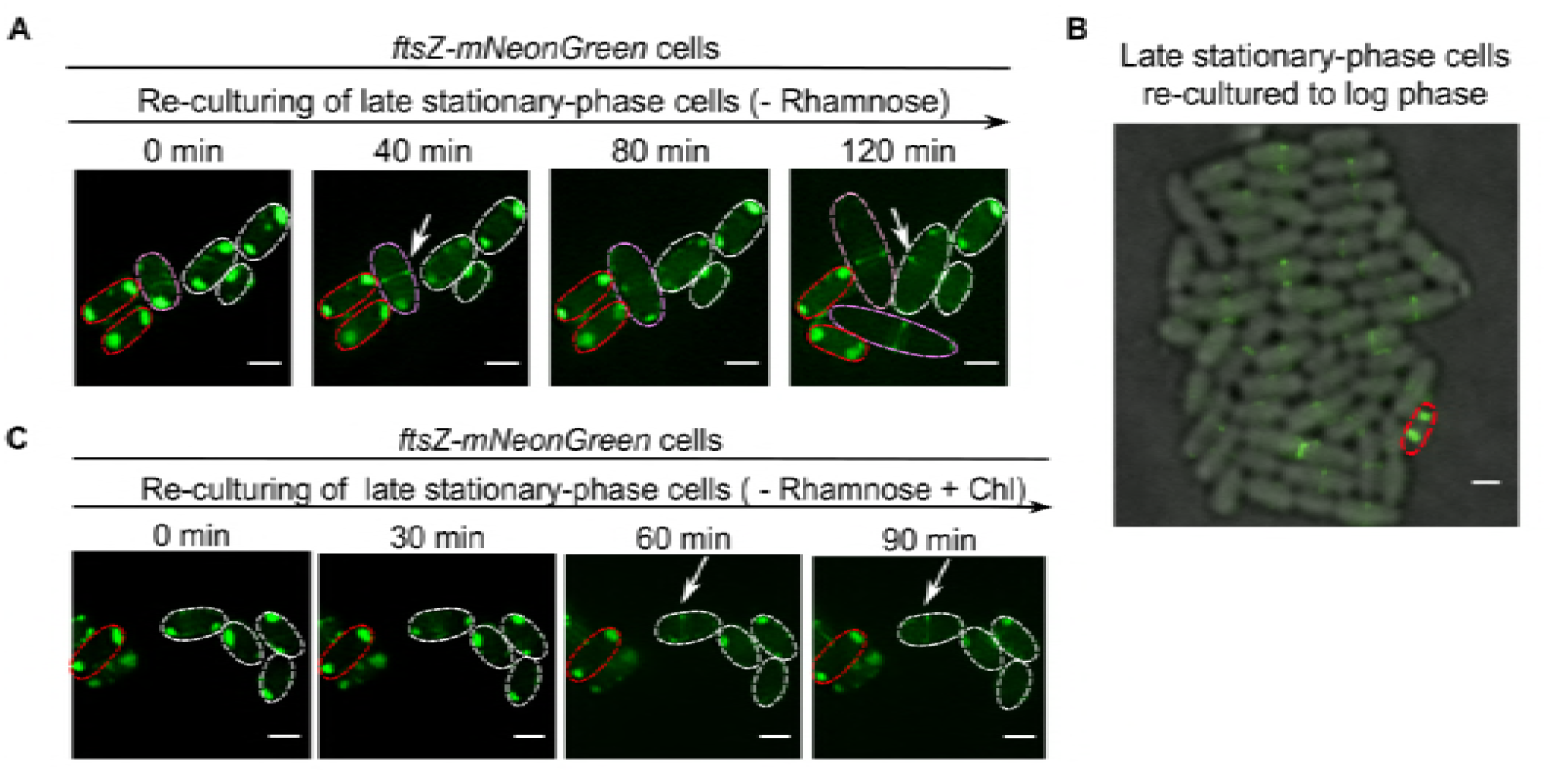
When the non-growing cells exit their regrowth lag, the cell-pole granules dissolve to release the FtsZ for re-functioning, but maintain unaltered otherwise. **(A)** Fluorescence microscopic images of re-cultured non-growing late stationary-phase *ftsZ-mNeonGreen* cells present in fresh LB medium lacking rhamnose, as obtained at the indicated time points. Note: one of the examined cells divided into two daughter cells at 120 min (circled by pink dashed lines). Scale bars, 1 μm. **(B)** Fluorescence microscopic images of the non-growing late stationary-phase *ftsZ-mNeonGreen* cells re-cultured to the log phase (OD_600_ ∼0.5) in liquid LB medium lacking rhamnose. Scale bar, 1 μm. **(C)** Fluorescence microscopic images of the non-growing late stationary-phase *ftsZ-mNeonGreen* cells re-cultured to the indicated time points in fresh LB medium that lacked rhamnose and contained the antibiotic chloramphenicol. Scale bars, 1 μm. (See also **Fig. S5**)

Worth of high attention, when the non-growing late stationary-phase *ftsZ-mNeonGreen* cells were recultured in fresh liquid medium lacking rhamnose to the log phase (with an OD_600_ of ∼0.5), we observed the maintenance of the cell-pole granules in an extremely small number of cells (as represented by the cell circled with red dashed lines in **Fig. 2B**), with all other cells being actively dividing. In our opinion, there is little doubt that such an inert cell, which is resistant to antibiotic killing (as to be shown in **Fig. 3F)**, should represent the long-searched and elusive persisters (Bigger, 1944).

**Figure 3.**
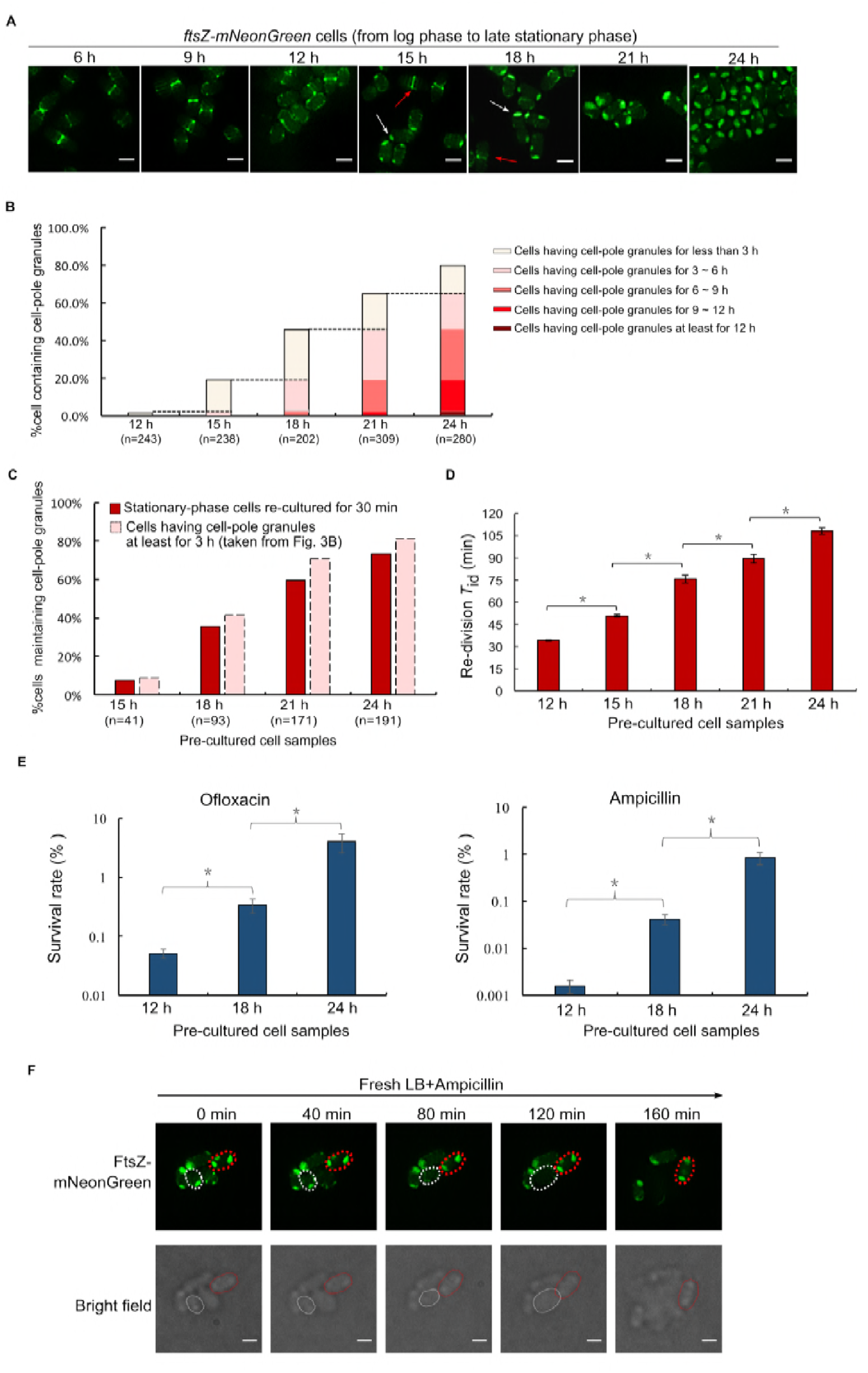
The regrowth-delay bodies are formed in a highly progressive in each individual cell and heterogeneous manner and in the cell population; the degree of their formation correlates with the duration of the regrowth lag time and the level of multidrug tolerance for a cell population. **(A)** Fluorescence microscopic images of *ftsZ-mNeonGreen* cells cultured to the indicated time points in LB medium containing 0.02% rhamnose (to induce the production of FtsZ-mNeonGreen). Scale bars, 1 μm. **(B)** Percentage of cells possessing regrowth-delay bodies within those cultured to the indicated time points (as shown in **A**). The values are shown in an accumulative manner (i.e., earlier values to be included in later values). **(C)** Percentages of cells maintaining their regrowth-delay bodies when the particular non-growing stationary-phase cell samples were re-cultured for 30 min in fresh medium containing chloramphenicol (red columns). Percentages of cells containing regrowth-delay bodies for more than 3 hours (pink columns) were directly taken from **Fig. 3B** to indicate their best fit matches as indicated by regression analysis. **(D)** Re-division *T*_id_ (the average initial doubling time) values of wild-type cells that were pre-cultured to the indicated time points. The *T*_id_ values were calculated based on the increase in cell numbers within the first 30 min of re-culturing (after diluting 40fold) in fresh medium at 37°C (for details, see Methods). (See also **Fig. S6**) **(E)** Survival rates of the indicated re-cultured non-growing stationary-phase wild-type cells that were treated with ofloxacin (5 μg/ml) or ampicillin (200 μg/ml) for 2 h (in fresh LB medium at 37°C). The survival rates were calculated according to the equation: [colony-forming units (CFU) of the antibiotic-treated cells] / [colony-forming units of the untreated cells] × 100. **(F)** Live-cell fluorescence (top) and bright field (bottom) microscopic images of the recultured non-growing late stationary-phase *ftsZ-mNeonGreen* cells in the fresh ampicillin-containing LB medium (at 37°C), as obtained at the indicated time points. One representative cell that exited (eventually became lysed) or maintained (unaltered) the regrowth lag is indicated by the white or red dashed circle, respectively. Scale bars, 1 μm. The symbol * in (**D**), (**E**) and (**C**) denotes a significant difference between the compared pair of samples (P-value <0.05, *t*-test). At least three biological replicates were analyzed in obtaining each value.

To assess whether such dissolution of the cell-pole granules could occur in the absence of any new synthesis of proteins in the cells, we repeated the above analysis by adding chloramphenicol, a ribosome-binding antibiotic that is known to inhibit protein synthesis in bacterial cells, to the fresh culture medium. Interestingly, we still observed an effective dissolution of the cell-pole granules in cells exiting the regrowth-lag and resuming growth (**Fig. 2C**, cells circled with white dashed lines), and we even occasionally observed the re-formation of Z-ring structures in certain cells (as indicated by the arrow). Of note, here the Z-ring structure would have to be formed mainly by using the FtsZ stored in and released from the cell-pole granules, but fluorescently labeled by a small amount of the incorporated FtsZ-mNeonGreen (Ma et al., 1996), also released from the granules. We again observed the cell-pole granules to be effectively retained in cells that remained in the regrowth-lag (**Fig. 2C**, exemplified by the cell circled with red dashed lines). In agreement with these live-cell imaging data, our immunoblotting analysis verified a time-dependent decrease of FtsZ in the insoluble pellet (**Fig. S6**, lanes 3, 6, and 9), with a corresponding increase of FtsZ in the soluble supernatant (lanes 2, 5 and 8) when the non-growing wild-type late stationary-phase cells were re-cultured in fresh LB (Lysogeny Broth) medium containing chloramphenicol.

Importantly, the results displayed in **Fig. 2** also revealed that the cell-pole granules present in different individual cells seem to exhibit a high degree of heterogeneity. Specifically, the cell-pole granules became totally dissolved, partially dissolved, or remained almost completely unaltered depending on the particular cell (clearly shown by the cells viewed at 120 min in **Fig. 2A** as well as those at 90 min in **Fig. 2C**). These data meanwhile suggest that an effective dissolution of the cell-pole granules is a prerequisite for a cell to end its regrowth-lag and resume growth, whereas the lack of their dissolution marks the maintenance of the non-growing persister state for a cell (as represented by the one shown in **Fig. 2B**).

### The cell-pole granules are formed in a highly heterogeneous manner in different individual cells and in a progressive manner in each cell

We next attempted to learn more about the nature of the manifested heterogeneity of the cell-pole granules (as shown in **Fig. 2**), by examining their formation process in the non-growing bacterial cells. For this purpose, we initially planned to employ a microfluidic chip device to monitor both the formation, during the non-growing phase, and the dissolution, during the regrowth phase, of the cell-pole granules in single *ftsZ-mNeonGreen* cells. Unfortunately, our efforts were unsuccessful, mainly because we were unable to set up a culturing condition under which the cell-pole granules were formed in the bacterial cells being placed in the available microfluidic system (likely due to a lack of high cell density or other unknown factors). Given this failed attempt, we then decided to address this issue by analyzing the cell population.

We started by conducting a qualitative live-cell imaging analysis to assess how cell-pole granules are formed in the non-growing *ftsZ*-*mNeonGreen* cells along the culturing process. The data, displayed in **Fig. 3A**, revealed that the formation of cell-pole granules appears to occur in a progressive manner in each individual cell, as the sizes of the cell-pole granules appeared to be different in different individual cells. Meanwhile, the data indicates a high heterogeneity in regards to the formation of cell-pole granules among the cell population. For instance, at 15 h of culturing, cell-pole granules are formed in some of the cells, while a small portion of other cells (indicated by the red arrow) were still dividing (i.e., with Z-ring structures remained visible).

Subsequently, we performed a quantitative live-cell imaging analysis to calculate the percentage of cells in which cell-pole granules were formed at the different culturing time points. As displayed in **Fig. 3B**, the percentage of cells containing the cell-pole granules clearly increased along the culturing process (data being shown in an accumulative manner at each time point). Together, these results indicate that the cell-pole granules are formed in a highly heterogeneous manner among the individual bacterial cells and in a progressive fashion in each individual cell.

### Bacterial cells containing more aged cell-pole granules stay in their regrowth-lag state for longer duration

To uncover the potential differences between the cell-pole granules present at different stationary-phase culturing time points, we measured the percentage of cells that retained their cell-pole granules (thus remaining in a regrowth-lag state) after being recultured in chloramphenicol-containing (and rhamnose-lacking) fresh medium for 30 min. This condition, being similar to that under which the 30 min imaging data shown in **Fig. 2C** was obtained, represents one under which cells whose cell-pole granules become fully dissolved (thus exiting the regrowth-lag state) could be effectively distinguished from the cells whose cell-pole granules remained largely unaltered (thus remaining in the regrowth-lag state).

The data, shown in **Fig. 3C** (red columns) clearly indicate that a higher percentage of cells remained in the regrowth-lag state when the non-growing cell samples were taken from a later stationary phase culturing point. A regression analysis revealed the best fit between the percentages of cells remaining in regrowth-lag state during reculturing and the percentages of cells whose cell-pole granules had existed for more than 3 hours during the stationary-phase (pink columns in **Figs. 3C**). These data suggest that for each individual bacterial cell, the more aged its cell-pole granules, the longer its duration of regrowth lag.

This correlation between the duration of the regrowth-lag and the age of the cell-pole granules was further demonstrated by comparing the average re-division initial doubling times (re-division *T*_id_) manifested by the cells that were taken from different stationary-phase culturing time points and recultured. Here, for each cell sample, the re-division *T*_id_ value was calculated based on its re-culturing growth curve (as displayed in **Fig. S7**), and reflects its duration of regrowth-lag. The data, presented in **Fig. 3D**, clearly reveal a higher re-division *T*_id_ value for a cell sample taken from a later culturing point in the stationary phase. Collectively, these results suggest that for each bacterial cell the duration of its regrowth-lag is apparently related to the status of its cell-pole granules. In light of this, we hereafter designate the cell-pole granule as the regrowth-delay body.

### Bacterial cells containing the regrowth-delay bodies are multidrug tolerant

We then assessed whether bacterial cells that contain regrowth-delay bodies are tolerant to multiple antibiotics, being a major feature attributed to persisters. To this end, we first compared the antibiotic tolerance capacity of the non-growing cells taken from different stationary-phase culturing time points. The data clearly show that the bacterial cells derived from a later culturing point, thus possessing a higher level of aged regrowth-delay bodies, exhibited a significantly higher level of tolerance towards the two examined antibiotics, either ofloxacin or ampicillin (**Fig. 3E**). More importantly, our live-cell imaging data provide direct evidence showing that bacterial cells retaining their regrowth-delay bodies would effectively survive the ampicillin treatment during the re-culturing process (as represented by the cell circled by red dashed lines in **Fig. 3F**). By contrast, cells having their regrowth-delay body dissolved would be efficiently killed (eventually lysed) under the same reculturing condition (as represented by the cell circled by while dashed lines in **Fig. 3F**).

Collectively, our observations, as shown in **Figs. 2** and **3**, strongly suggest that the regrowth-delay bodies serve as effectively markers for the non-growing and antibiotic-tolerant bacterial persisters. It follows that the presence of regrowth-delay bodies would help us to efficiently identify the tiny subpopulation of persisters present in a large population of actively growing bacterial cell (as exemplified by the data shown in **Fig. 2B**). Our data meanwhile implicate that persister cells are in different depth of persistence depending on the age of their regrowth-delay bodies.

### The formation of regrowth-delay bodies selectively sequesters multiple key proteins

We next attempted to characterize the composition of the regrowth-delay bodies to learn more about the properties of bacterial persisters. For this goal, we first tried to identify the proteins that are photo-crosslinked to multiple pBpa variants of the FtsZ protein in the non-growing late stationary phase cells. Specifically, we purified the photo-crosslinked products of five FtsZ variants with pBpa introduced at residue position 140 (lane 8 in **Fig. S1B**), 47, 51, 61 or 166 (lanes 10, 12, 16 or 4, respectively, in **Fig. S5D**), each representing a different pattern of photo-crosslinked products, by affinity chromatography via the Avi tag fused to the FtsZ protein. The proteins were identified via mass spectrometry analysis and are listed in **Fig. S8A**.

In light that intact regrowth-delay bodies were present in the pellet fraction (as shown in **Fig. S4C**), we also performed mass spectrometry analysis on the collected pellet of the lysed wild-type *E. coli* cells, with the major proteins identified being also listed in **Fig. S8A**. A functional annotation of these identified proteins revealed their key roles in cell growth (such as translation and transcription) and division, which in part explains why their sequestering in the regrowth-delay bodies could keep the bacterial cells in a non-growing state. Of note, some of the proteins (colored blue in **Fig. S8A**) were identified by both mass spectrometry analyses.

We subsequently performed experiments to verify the presence of some of these identified proteins (other than FtsZ) in the regrowth-delay bodies. We first confirmed by live-cell imaging analysis that ZapC and FtsA (each being fused to mNeonGreen), two additional cell division proteins identified, were both clearly detected in the regrowth-delay bodies as present in the non-growing late stationary-phase cells, though in the Z-ring structure in actively dividing log-phase cells (**Fig. 4A**). We also demonstrated that FtsA (as fused to the red fluorescent protein mCherry) co-localizes with FtsZ (as fused to mNeonGreen) in the regrowth-delay bodies present either in living cells or in the lysates (**Fig. S8B**). By contrast, FtsL and ZapA, two non-identified cell division proteins, were neither detected in the regrowth-delay bodies, but evenly distributed in the cytosol in the non-growing late stationary-phase cells, while clearly detected in the Z-ring structures in actively dividing log-phase cells (**Fig. S8C**).

**Figure 4.**
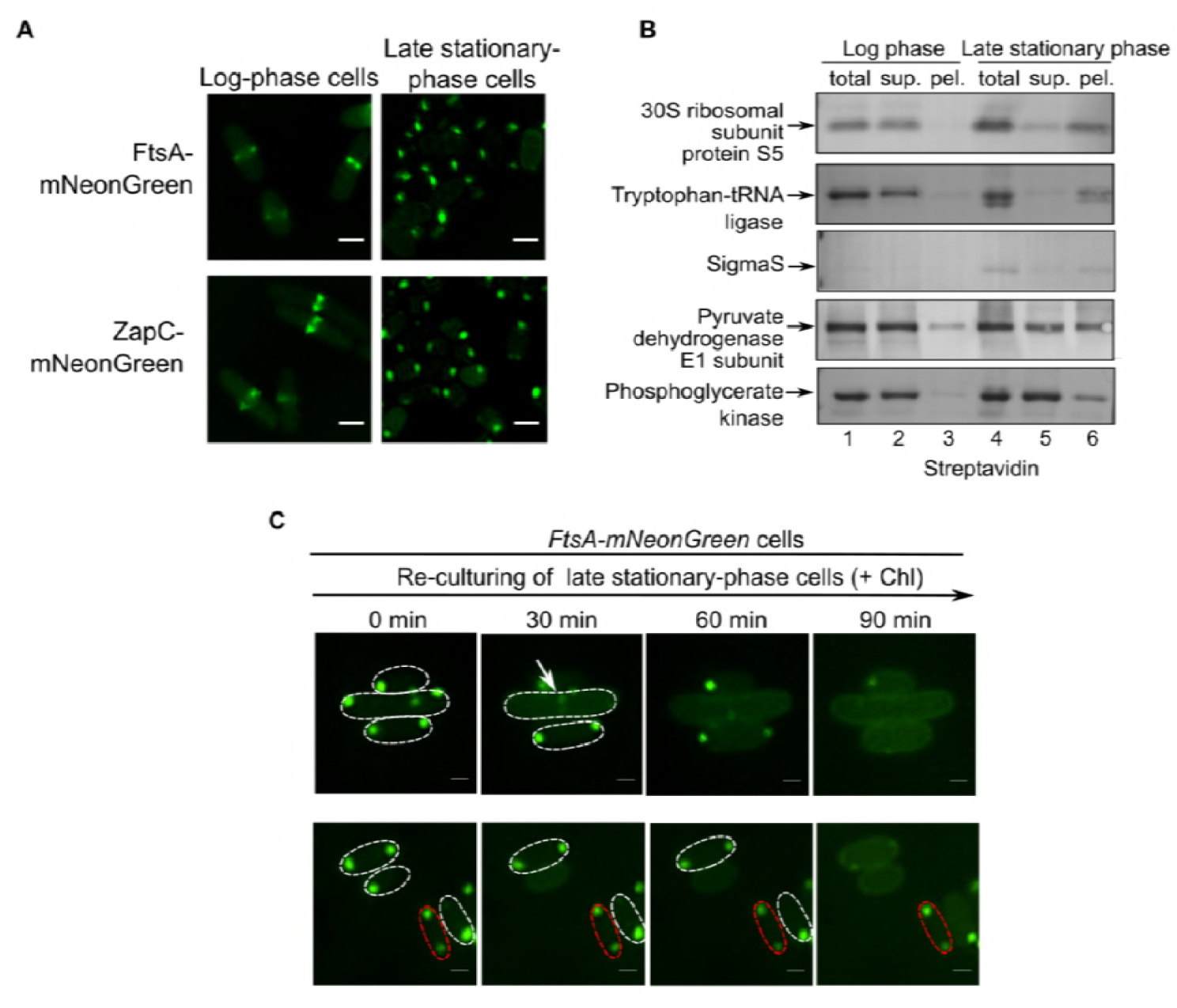
Regrowth-delay bodies selectively sequester multiple other key proteins that are released to re-function when cells exit their regrowth lag and resume growth. **(A)** Fluorescence microscopic images of actively dividing log-phase (left) and nongrowing late stationary-phase (right) *E. coll* cells in which mNeonGreen-fused cell division FtsA or ZapC (both being identified in the regrowth-delay bodies by mass spectrometry analysis, as shown in **Fig. S7A**) was expressed from a plasmid under the control of a constitutive promoter. Scale bars, 1 μm. (See also **Figs. S7A** and **S7B**) **(B)** Blotting results to analyze the indicated Avi-tagged proteins (all being identified in the regrowth-delay bodies by mass spectrometry analysis, as shown in **Fig. S7A**, and each being expressed from a plasmid under the control of a constitutive promoter) in the indicated lysate fractions of actively dividing log-phase or non-growing late stationary-phase wild-type cells, probed with streptavidin-AP. **(C)** Fluorescence microscopic images of two fields of re-cultured non-growing late stationary-phase cells, in which FtsA-mNeonGreen was expressed from a plasmid under the control of a constitutive promoter, in fresh medium containing chloramphenicol, obtained at the indicated time points. Scale bars, 1 μm.

We then verified the apparent presence of five more identified proteins. In particular, they, each being expressed as an Avi-tagged form and under the control of a constitutive promoter, were detected to a significant degree in the insoluble pellet fraction of lysates of the non-growing late stationary-phase cells, though largely present in the supernatant of lysates of actively dividing log-phase cells (**Fig. 4B**). Interestingly, among these five proteins, the three that were known to be essential for cell growth (i.e., ribosomal protein S5, tryptophan-tRNA ligase, and transcriptional factor sigmaS) were almost fully detected in the pellet fraction (**Fig. 4B**). Of note, the sigmaS protein is known to be degraded in actively dividing log-phase cells and accumulates only in non-growing stationary-phase cells (Zhou and Gottesman, 1998). Taken together, these protein characterization and verification studies clearly suggest that the regrowth-delay bodies sequester multiple important proteins that function in cell growth and division, which in turn conceivably keep the cells in the non-growing persister state.

Additionally, we demonstrated by performing live-cell imaging analysis that similar to FtsZ, FtsA was also reutilized in cells exiting the regrowth-lag and resuming growth (**Fig. 4C**). Specifically, the FtsA protein (fused to mNeonGreen) either reappeared in the Z-ring structure (shown by the arrow in **Fig. 4C**) of cells that were in the process of re-dividing or in the inner membrane of cells whose regrowth-delay bodies were dissolved but not yet dividing as reported before (Pichoff and Lutkenhaus, 2005) when the non-growing late stationary-phase cells were recultured in fresh medium containing chloramphenicol. Similarly, the FtsA protein was retained in the regrowth-delay bodies for cells remaining in the non-growing regrowth-lag state (**Fig. 4C**, represented by the cell circled by red dashed lines). These results once again demonstrated that the proteins sequestered in the regrowth-delay bodies are released to resume their functions in cells exiting the regrowth-lag state and resuming growth.

### Mutant bacterial cells with a reduced formation of regrowth-delay bodies exhibit a shorter duration of regrowth lag and a lower tolerance to antibiotics

To further examine the relationship between regrowth-delay body formation and regrowth lag time or antibiotic tolerance, we then attempted to generate mutant bacterial cells in which the formation of regrowth-delay bodies would be significantly reduced. Towards this goal, we referred to the list of proteins identified in the regrowth-delay bodies (as shown in **Fig. S8A**) and realized the presence of multiple subunits of the respiratory chain complexes. Furthermore, our live-cell imaging analysis (**Fig. S4A**) showed an apparent association of the regrowth-delay bodies with the inner membrane, where the respiratory chain complexes are located. In light of these observations, we then performed gene knockdown (or gene knockout) experiments to decrease or remove certain subunits of the respiratory chain complexes and analyzed whether the regrowth-delay body formation in the bacterial cells was significantly reduced. In particular, the *nuoA* gene (encoding a subunit of respiratory chain complex I) or the *sdhC* gene (encoding a subunit of respiratory chain complex II) in the *ftsZ-mNeonGreen* cells was subjected to knockdown manipulation using the CRISPRi technology (Luo et al., 2015).

Our live-cell imaging analysis (**Fig. 5A**, left panel) demonstrated that the regrowth-delay body formation was significantly reduced in the non-growing late stationary-phase *nuoA*-knockdown cells and barely occurred in the *sdhC*-knockdown cells. In agreement with these imaging results, our immunoblotting analysis confirmed a significantly reduced amount of the endogenous FtsZ in the insoluble lysate pellet fraction of these cells, instead, much appeared in the soluble supernatant fraction (**Fig. 5A**, right panel). We observed similar reduction in regrowth-delay body formation (shown in **Fig. S9A**) for the bacterial cells in which the *nouAB* (genes encoding two subunits of respiratory chain complex I) or *sdhCDAB* (genes encoding all the four subunits of the respiratory chain complex II) were knocked out. Taken together, these observations indicate that the respiratory chain complexes somehow do play an important role for the formation of regrowth-delay bodies.

**Figure 5.**
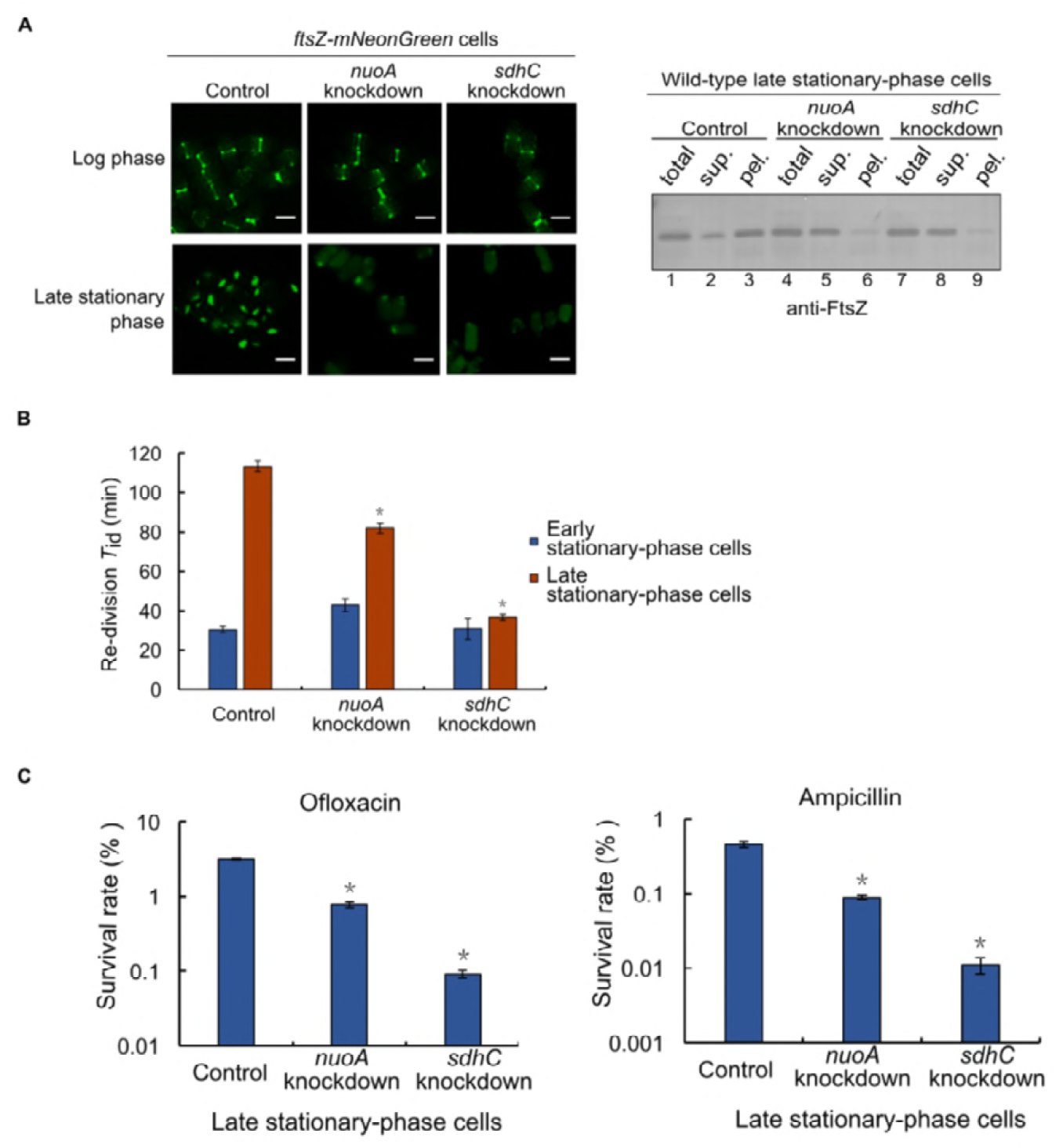
Mutant bacterial cells with a reduced formation of regrowth-delay bodies exhibit a shorter duration of regrowth lag as well as a lower tolerance to antibiotics. **(A)** Fluorescence microscopic images of actively dividing log-phase or non-growing late stationary-phase *ftsZ-mNeonGreen* cells having a knockdown of either the *nuoA* or the *sdhC* gene. Cells expressing a non-targeting CRISPR RNA were analyzed as the control, scale bars, 1 μm (left panel); The immunoblotting results for detecting FtsZ in the indicated cell lysate fractions, as probed with anti-FtsZ antibodies (right panel). (See also **Fig. S8A**) **(B)** Re-division *T_id_* values of early (blue bars; cultured to 12 h) or late (red bars; cultured to 24 h) non-growing stationary-phase cells of the indicated gene-knockdown strain. Here wild-type cells in which a non-targeting crRNA was expressed from a plasmid were analyzed as the control. (See also **Fig. S8B**) **(C)** Survival rates of the non-growing late stationary-phase wild-type (control), *nuoA-* knockdown or sdhC-knockdown cells that were re-cultured in fresh medium after being treated with ofloxacin (5 μg/ml) or ampicillin (200 μg/ml). The survival rates were calculated according to the equation: [CFU of the antibiotic-treated cells] / [CFU of the untreated cells] × 100. The symbol * in (**B**) and (**C**) denotes a significant difference between the compared pair of samples (P-value <0.05, *t*-test). At least three biological replicates were analyzed for obtaining each value.

In agreement with our hypothesis, we observed that the re-division *T*_id_ value of either the *nuoA* or *sdhC* knockdown cells was significantly lower in comparison with that of the control cells (**Figs. 5B**). Additionally, the re-division *T*_id_ values became comparable for the early and late stationary-phase *sdhC* knockdown cells, unlike those for the control cells (**Fig. 5B**). Consistently, the survival rates of these non-growing late stationary-phase cells became significantly lower than those of the control cells after being treated with an antibiotic, ofloxacin or ampicillin (**Fig. 5C**). These observations on the gene knockdown cells further strengthened our conclusion that bacterial cells exhibit a prominent regrowth lag and antibiotic tolerance due to the formation of regrowth-delay bodies.

### Regrowth-delay body formation occurs in pathogenic bacteria and also correlates to the regrowth-lag and multidrug tolerance

We subsequently demonstrated the formation of regrowth-delay bodies in such pathogenic bacteria as *Salmonella* Typhimurium and *Shigella flexneri*, which respectively cause gastroenteritis and diarrhea in humans (Graham, 2002; Jennison and Verma, 2004). In particular, we observed a similar time-dependent appearance of the endogenous FtsZ in the lysate pellet of non-growing late stationary-phase cells for either *Salmonella* Typhimurium SL1344 or *Shigella flexneri* serotype 2a 2457T (**Fig. 6A**). For each strain, we then observed a similar correlation between a higher degree of regrowth-delay body formation and a longer regrowth lag time (**Fig. 6B** and **Fig. S10**) or a higher level of antibiotic tolerance (**Fig. 6C**). These observations again indicate that regrowth-delay body formation likely attributes to the regrowth lag and antibiotic tolerance in bacterial persister cells.

**Figure 6.**
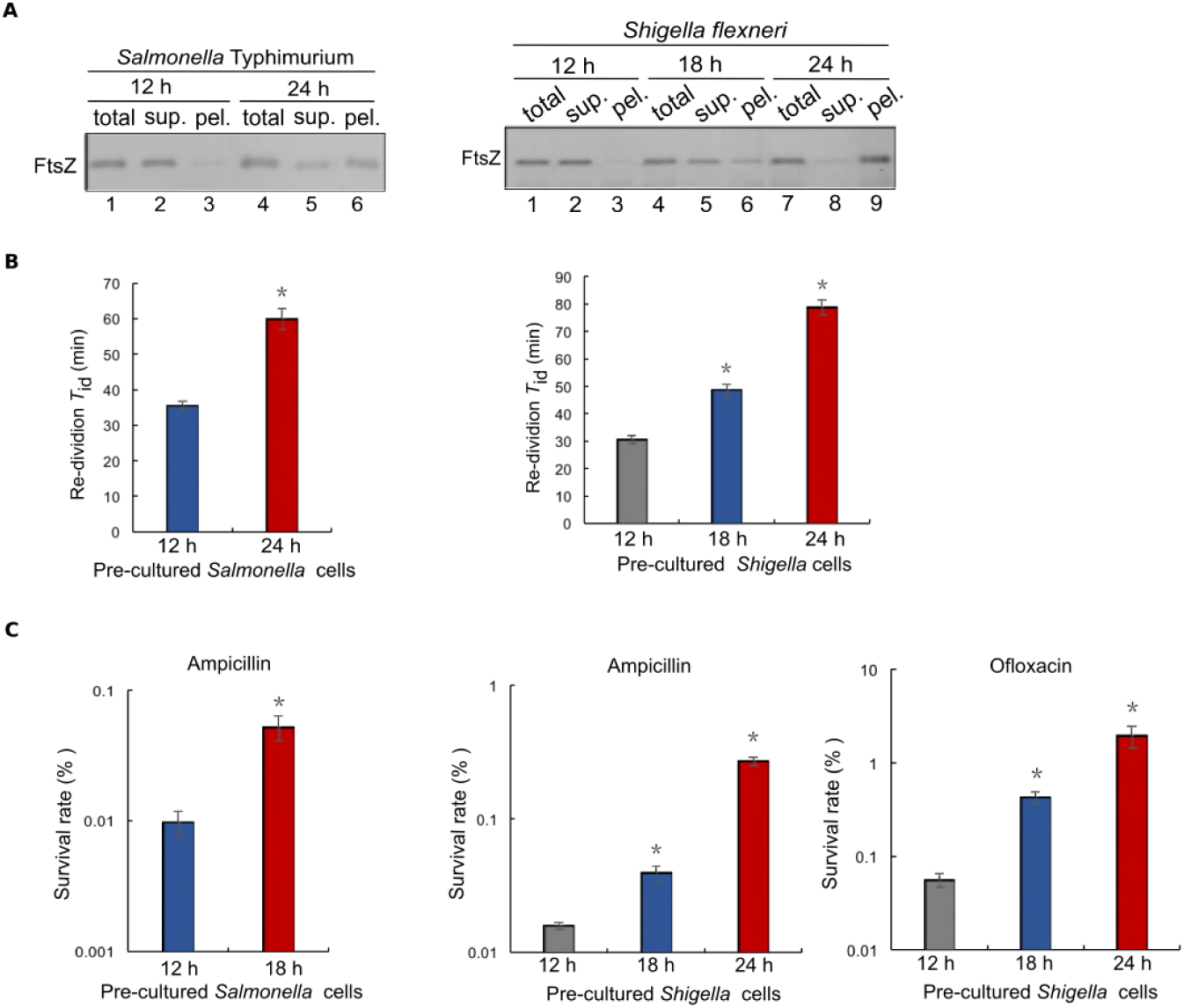
Regrowth-delay bodies are also formed in the non-growing late stationary-phase cells of the pathogenic bacteria *Salmonella* Typhimurium SL1344 and *Shigella flexneri* serotype 2a 2457T. **(A)** Immunoblotting results for the detection of FtsZ in the indicated cell lysate fractions of the non-growing stationary-phase *Salmonella* Typhimurium or *Shigella flexneri* cells taken at the indicated time points, probed with antibodies against the *E. coli* FtsZ protein. **(B)** Re-division *T*_id_ values of the non-growing *Salmonella* Typhimurium or *Shigella flexneri* cells that were pre-cultured to the indicated time points of the stationary-phase before being re-cultured in fresh LB medium. (See also **Fig. S9**) **(C)** Survival rates of the non-growing *Salmonella* Typhimurium or *Shigella flexneri* cells that were pre-cultured to the indicated time points of the stationary-phase before being re-cultured in fresh LB medium after being treated with the indicated antibiotics for 2 h. The symbol * in (**B**) and (**C**) denotes a significant difference between the compared pair of samples (*P*-value <0.05, *t*-test). At least three biological replicates were analyzed for obtaining each value.

## DISCUSSION

Here, we reported our accidental discovery of a hitherto unreported bacterial subcellular structure that we designated as the regrowth-delay body. In retrospect, we made this revelation as a result of our initial *in vivo* protein photo-crosslinking and subsequent live-cell imaging analyses on the unique FtsZ protein, not only with actively dividing cells (as have been extensively examined by others), but also with the nondividing/non-growing cells (as have been rarely examined by others). We provided ample evidence to support our conclusion that the regrowth-delay bodies are formed by sequestering multiple key cellular proteins, which in turn enable bacterial cells to enter a persister state, which exhibits not only a regrowth lag but also a multidrug tolerance.

Regrowth-delay body represents a distinctive subcellular structure that allows the tiny subpopulation of persisters to be effectively identified in a large population of actively growing cells, a prerequisite for elucidating their physiological properties. Meanwhile, our demonstration that regrowth-delay bodies sequesters multiple key cellular proteins provides key mechanistic insights for explaining why persisters are able to maintain in a non-growing dormant state for an extended period of time, being an outstanding unresolved puzzle in microbiology.

Importantly, our findings imply that a bacterial persister is actually in a particular depth of persistence, as determined by the status of its regrowth-delay bodies. In other words, a persister whose regrowth-delay bodies are to be dissolved rather effectively is in a shallow persistent state, thus to exhibit a relatively short regrowth lag whenever as an optimal growth condition becomes available. Conversely, a persister whose regrowth-delay bodies are to be maintained for an extended period of time even when an optimal growth condition becomes available is in a deep persistent state. According to this, a conventional multidrug-tolerant persister represents a bacterial cell that is in deep persistence, or a metabolically inactive dormant state.

Having cells in different depths of persistence would conceivably allow certain number of persister cells to survive under any harmful condition. This explains how the formation of regrowth-delay bodies would provide an effective bet-hedging strategy for a bacterial species to maximize its possibility of survival in the highly unpredictable natural environment (Kell et al., 2015; Maisonneuve and Gerdes, 2014; Veening et al., 2008). In a sense, the regrowth-delay bodies in a persister cell function as the biological timer that determines the particular duration of the regrowth lag for the non-growing bacterial cell to resume growth

Our revelations also explain why the formation of persisters has long been viewed as a stochastic or heterogeneous phenomenon occurring in the bacterial cell populations (Allison et al., 2011; Gefen and Balaban, 2009; Amato and Brynildsen, 2015; Dhar and McKinney, 2007). This is mainly due to the high heterogeneity of regrowth-delay body formation in different individual cells as well as the progressive nature of their formation in each single cell. Because of this, a bacterial cell sample taken from different culturing point would be highly heterogeneous in regards of the status of the cellular regrowth-delay bodies or depth of persistence in different cells. It follows that the duration of the regrowth lag, the level of drug tolerance, as well as the percentage of cells defined as persisters (by measuring the number of colony-forming units after treating an antibiotic) in the cell population, would most likely appear as inconsistent or stochastic values even in repeating experiments.

One difficulty in studying the persister cells is to unequivocally identify them, as they usually exist in extremely small numbers in a cell population that are actively growing (Balaban et al., 2013). The presence of the distinctive regrowth-delay bodies in persisters would prove to be greatly helpful in overcoming this difficulty (as exemplified by the data shown in **Fig. 2C**). This meanwhile may allow us to conduct single cell biochemistry and cell biology studies on persisters, including a characterization of the transcriptomes, proteome and metabolome (Kell et al., 2015; Taniguchi et al., 2010).

In light of our findings described here, the “viable but non-culturable” bacteria, which is known to evade the conventional culture-based microbiological detection (Pinto et al., 2015), may represent persister cells whose regrowth-delay bodies could not effectively dissolve under the commonly applied culturing conditions. After we learn more about the conditions that will effectively promote the dissolution of regrowth-delay bodies, we may be able to make these bacterial cells culturable under particular conditions. By the same token, in clinics, we might be able to find ways to eradicate the multidrug tolerant recalcitrant pathogen persisters by promoting the dissolution of their regrowth-delay bodies in conjunction with an antibiotic treatment.

However, many questions remain unanswered concerning the biology of regrowth-delay bodies, as a new subcellular structure marking the non-growing persister bacterial cells. First, how are the components in the regrowth-delay bodies organized (to be revealed likely by high resolution electron microscopic analysis)? Second, what are the key signaling molecules that trigger their formation, and how are such signals sensed by cells? Third, how are the specifically sequestered proteins selected? Fourth, what signals trigger the regrowth-delay bodies to dissolve? Finally, do structures similar to regrowth-delay bodies exist in eukaryotes, especially those living as single-cell forms?

## Author Contributions

Jiayu Yu and Yang Liu designed and performed the major experiments, analyzed the data, and drafted the manuscript. Huijia Yin designed and performed part of the experiments. Prof. Zengyi Chang supervised the entirety of the study.

## Acknowledgments

We thank Prof. Harold Erickson (Duke University, USA) for providing us the pJSB100 plasmid, and Prof. Peter Schultz (The Scripps Research Institute, USA) for providing us with the plasmids that carry the genes encoding the orthogonal tRNA and orthogonal amino acyl-tRNA for pBpa incorporation. We thank Keio Collections for providing us the wild-type *E. coli* strain. We thank Dr. Xiaoyun Liu (Peking University, China) for providing the *Salmonella* Typhimurium and *Shigella flexneri* strains. We thank the Core Facilities at the School of Life Sciences, Peking University, for assistance using the structured illumination microscope (SIM), and we are grateful to Dr. Chunyan Shan and Dr. Xiaochen Li for assisting us with the fluorescence microscopic imaging analysis. We thank Dr. Wen Zhou at the Mass Spectrometry Facility of the National Center for Protein Sciences at Peking University for assistance on the mass spectrometry analysis. We thank Prof. Chong Liu from Zhejiang University and Prof. Xinmiao Fu from Fujian Normal University for useful discussions. This work was supported by funds from the National Natural Science Foundation of China (No. 31670775 and 31470766 to ZYC), the National Basic Research Program of China (No. 2012CB917300 to ZYC), and the Qidong-SLS Innovation Fund. We declare that we have no conflicts of interest related to this work.

## Supplemental figures

**Figure S1.**
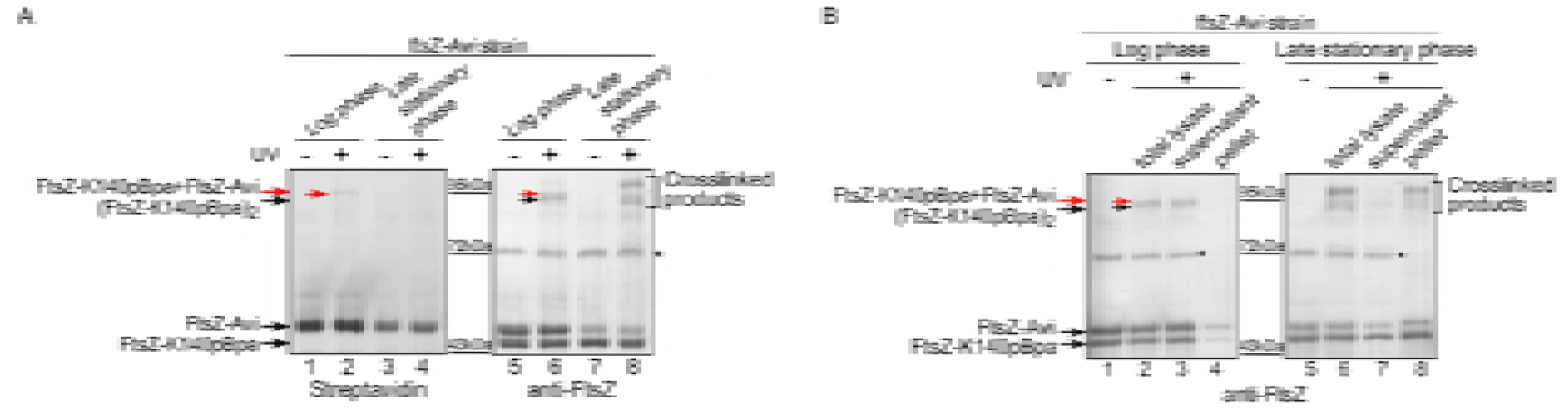
The cell division protein FtsZ exists as a self-assembled oligomer in actively dividing log-phase cells but as unassembled and insoluble form in nongrowing late stationary-phase *E. coli* cells. **(A)** Blotting results for the detection of photo-crosslinked products of the FtsZ-K140pBpa variant in the actively dividing log-phase and the non-growing late stationary-phase *ftsZ-Avi* cells exposed to UV light, as probed with streptavidin-alkaline phosphate conjugate (left part) or antibodies against FtsZ (right part). The asterisk indicates a non-specific protein band detected when probed with the anti-FtsZ antibodies. **(B)** Immunoblotting results for the detection of photo-crosslinked products of the FtsZ-K140pBpa variant, as well as the free FtsZ monomers, in the supernatant (sup.) and pellet (pel.) fractions of the actively dividing log-phase or non-growing late stationary-phase *ftsZ-Avi* cells, as probed with antibodies against FtsZ. Positions of the FtsZ monomers and photo-crosslinked dimers are shown on the left of the gels, positions of the molecular weight markers are shown in the middle of the gels (in both **A** and **B**). Here, to verify the reported *in vitro* assembly pattern of FtsZ protofilaments in *E. coli* cells, we performed *in vivo* protein photo-crosslinking analysis by replacing the amino acid residue K140, located at the longitudinal interface of the FtsZ protofilament, with the unnatural amino acid pBpa. This FtsZ-K140pBpa variant, which we demonstrated to be able to support cell division in the absence of wild type FtsZ, was then heterologously expressed in a strain whose own genomic *ftsZ* gene was modified to encode an Avi-tagged FtsZ variant (the Avi tag could be specifically probed with streptavidin). As shown by the blotting results displayed here in **Fig. S1A**, the FtsZ dimers were formed either between the FtsZ-K140pBpa and FtsZ-Avi monomers (thus detectable not only by streptavidin AP conjugate but also by antibodies against FtsZ; red arrows) or between two FtsZ-K140pBpa monomers (only detectable by antibodies against FtsZ; black arrows) in actively dividing log-phase cells (lanes 2 and 6). These observations confirmed the location of residue K140 at a self-assembling interface and that FtsZ assembles into homo-oligomers in actively dividing cells. By contrast, in non-growing late stationary-phase cells, the photo-crosslinked FtsZ dimers became no longer detectable (Fig. S1A, lanes 4 and 8), instead, multiple photo-crosslinked products between FtsZ-K140pBpa and other proteins were readily detected (lane 8). These results seem to indicate that residue K140 now mediate interactions with multiple other proteins in the non-growing bacterial cells.

**Figure S2.**
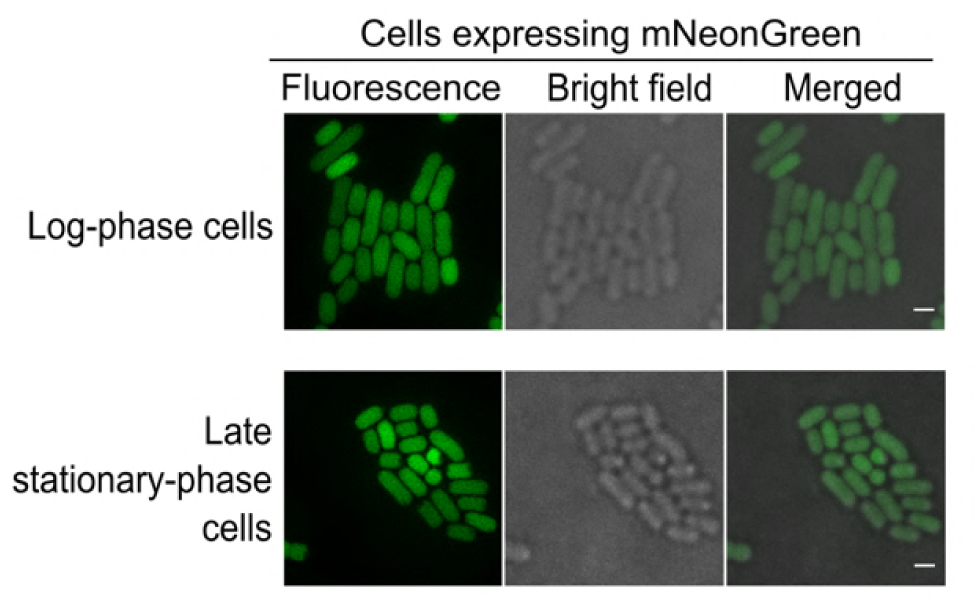
Fluorescence and bright-field microscopic images of the actively dividing log-phase (top) and the non-growing late stationary-phase (bottom) *E. coli* cells in which the green fluorescent protein mNeonGreen (without being fused to FtsZ) was heterologously expressed. Scale bars, 1 μm.

**Figure S3.**
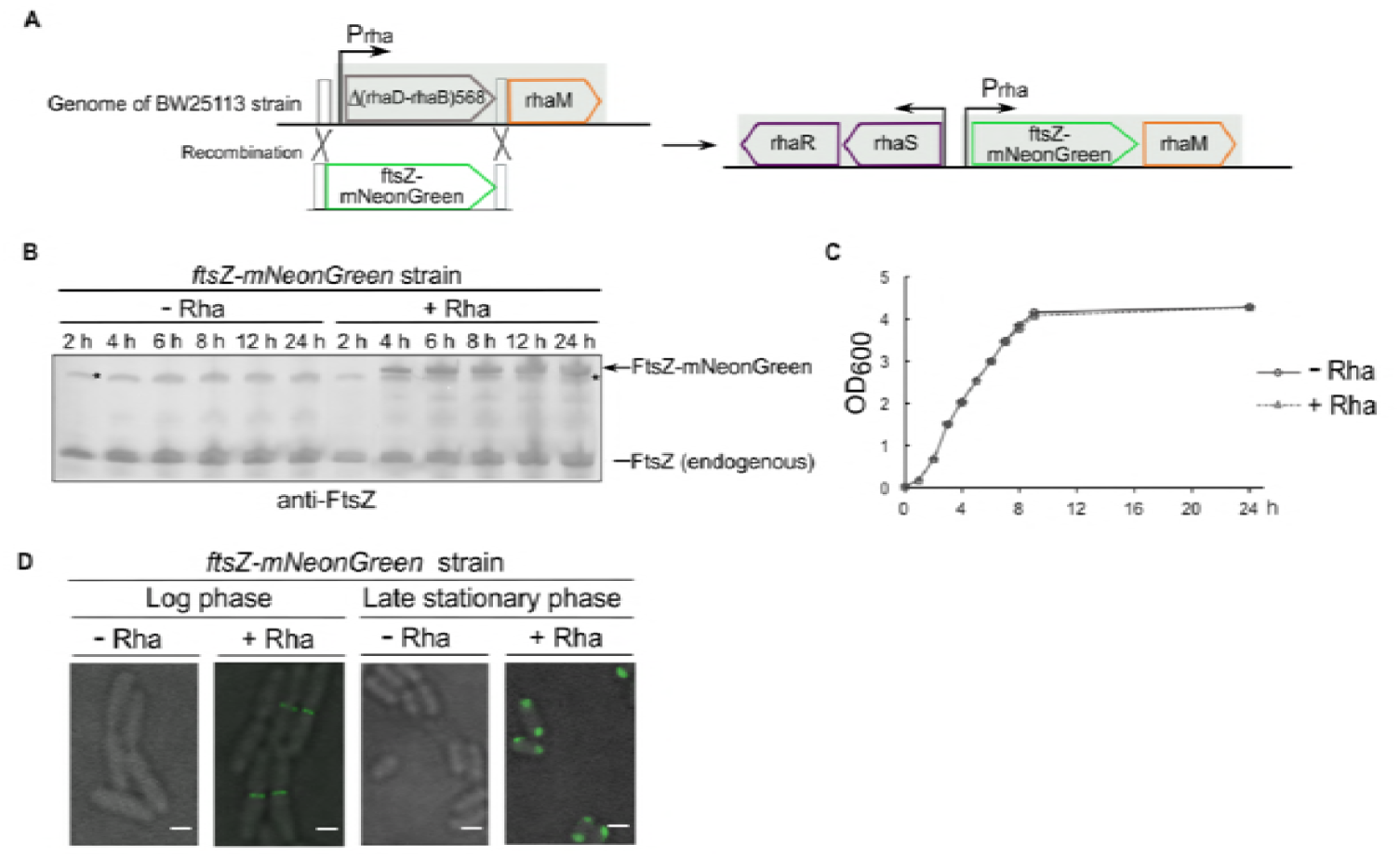
Construction and verification of the *ftsZ-mNeonGreen* strain. **(A)** The *ftsZ-mNeonGreen* strain was constructed by replacing part of the rhamnose operon by the *ftsZ-mNeonGreen* gene (green outline) in the *E. coli* genome. The transcription initiation sites and directions of transcriptions are both indicated by the arrows (top panel). Immunoblotting results for detecting the FtsZ-mNeonGreen protein expressed in the *ftsZ-mNeonGreen* strain as cultured in the presence (+Rha) or absence (-Rha) of rhamnose (0.02%) to the indicated time points, as probed with antibodies against FtsZ; positions of the two forms of FtsZ are indicated on the right. Asterisk indicates a non-specific protein band (bottom left panel). **(B)** Growth curves of the *ftsZ-mNeonGreen* strain cultured in the presence (+Rha) or absence (-Rha) of rhamnose (0.02%), as prepared by measuring the OD_600_ values at the indicated time points (bottom right panel). **(C)** Immunoblotting results for detecting the FtsZ-mNeonGreen protein expressed in the *ftsZ-mNeonGreen* strain cultured in the presence (+Rha) or absence (-Rha) of FtsZ. Positions of the two forms of FtsZ are indicated on the right. Asterisk indicates a nonspecific band. **(D)** Bright field and fluorescence microscopic images of the log-phase or late stationary-phase *ftsZ-mNeonGreen* cells cultured in LB media with (+Rha) or without (-Rha) the addition of rhamnose. Scale bars, 1 μm.

**Figure S4.**
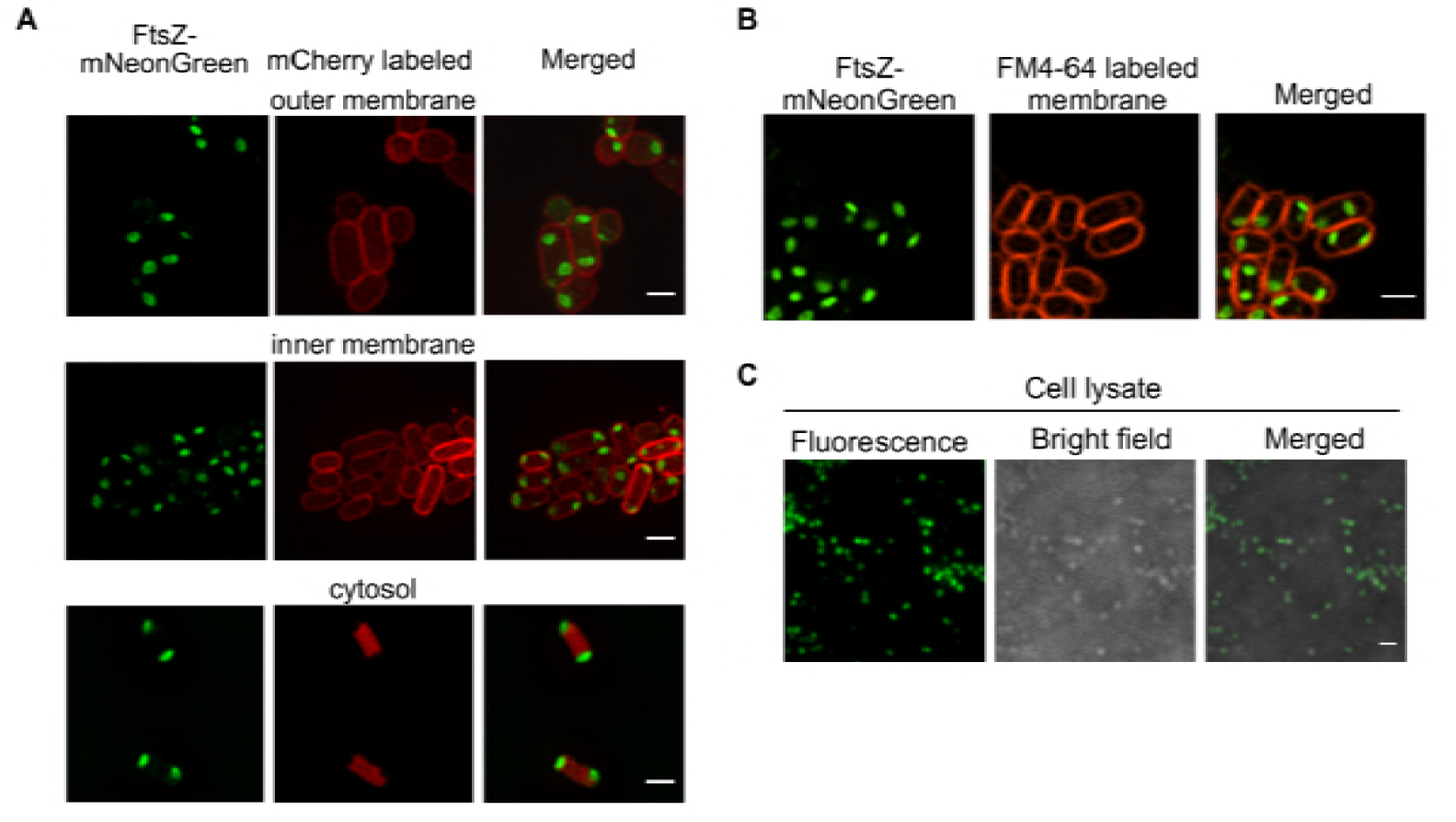
The regrowth-delay bodies exist as a compact subcellular structure in the cytosol and seem to be associated with the inner membrane but not surrounded by any membrane component. **(A)** Fluorescence microscopic images of the non-growing late stationary-phase *ftsZ-mNeonGreen* cells whose outer membrane (top), inner membrane (middle) or cytosol (bottom) was separately labeled with OmpA-fused mCherry, NlpA anchoring peptide-fused mCherry or unfused mCherry, respectively. Scale bars, 1μm. **(B)** Fluorescence microscopic images of the non-growing late stationary-phase *ftsZ-mNeonGreen* cells stained with the membrane specific FM4-64 dye. Scale bars, 1 μm. **(C)** Fluorescence and bright field microscopic images of the regrowth-delay bodies detected in the lysates of non-growing late stationary-phase (cultured to 24 h) *ftsZ-mNeonGreen* cells. Scale bars, 1 μm.

**Figure S5.**
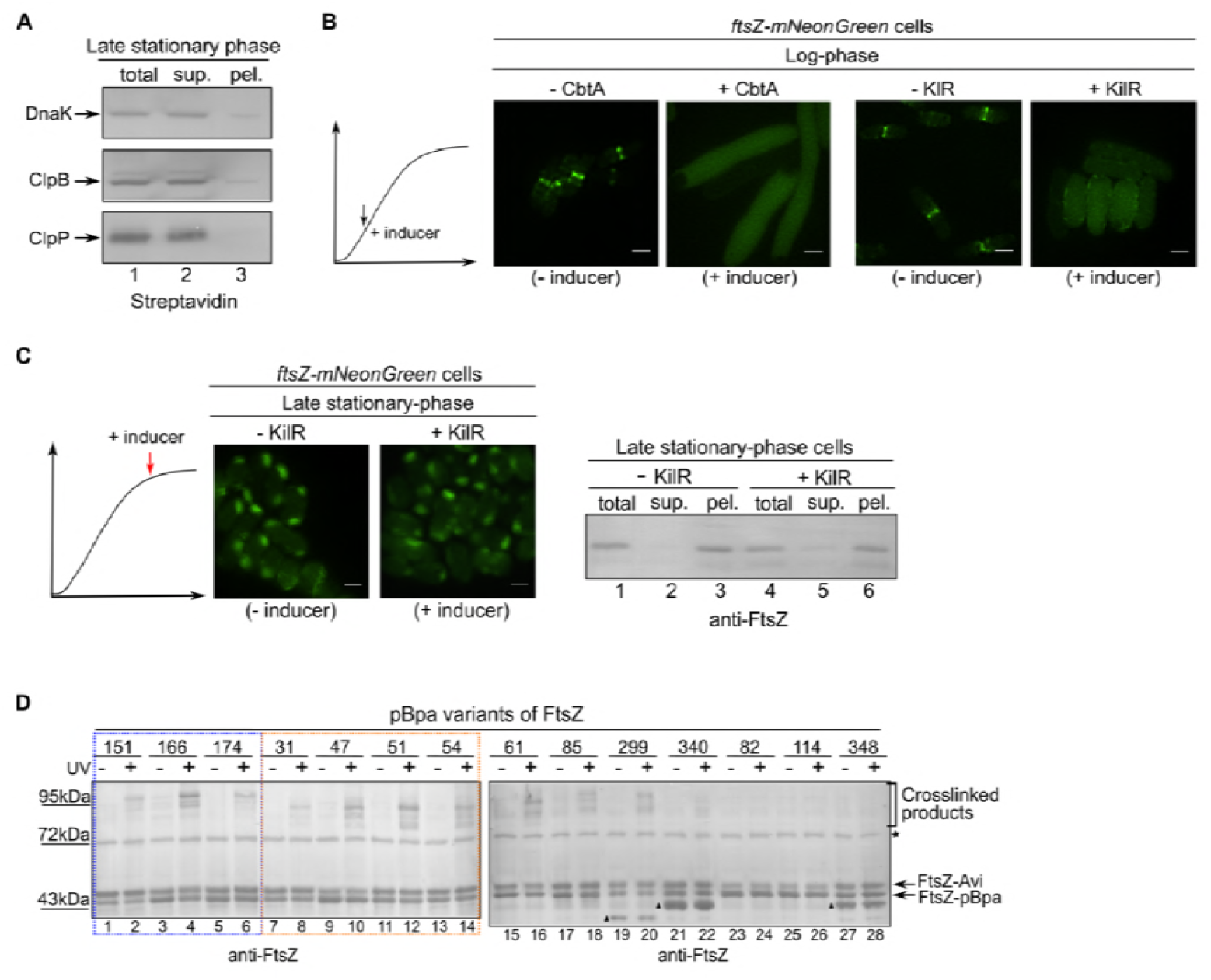
Expression of the inhibitor protein CbtA or KilR prevents FtsZ to assemble into the Z-ring structure in log-phase *ftsZ-mNeonGreen cells*. However, expression of KilR (in contrast to CbtA, as shown in Fig. 1D) does not prevent FtsZ to enter the regrowth-delay bodies. **(A)** Blotting results for the detection of DnaK-Avi, ClpB-Avi or ClpP-Avi protein in the indicated fractions of non-growing late stationary-phase *E. coli* cells, probed with the streptavidin-AP conjugate (against the Avi tag). **(B)** Fluorescence microscopic images of log-phase *ftsZ-mNeonGreen* cells in which the expression of the CbtA or KilR inhibitor protein was induced. Scale bars, 1 μm. **(C)** Fluorescence microscopic images of the non-growing late stationary-phase *ftsZ-mNeonGreen* cells in which the expression of KilR was induced (left panel) and the corresponding immunoblotting results for the detection of the indicated cell lysate fractions, probed with antibodies against FtsZ (right panel). Scale bars, 1 μm.

**Figure S6.**
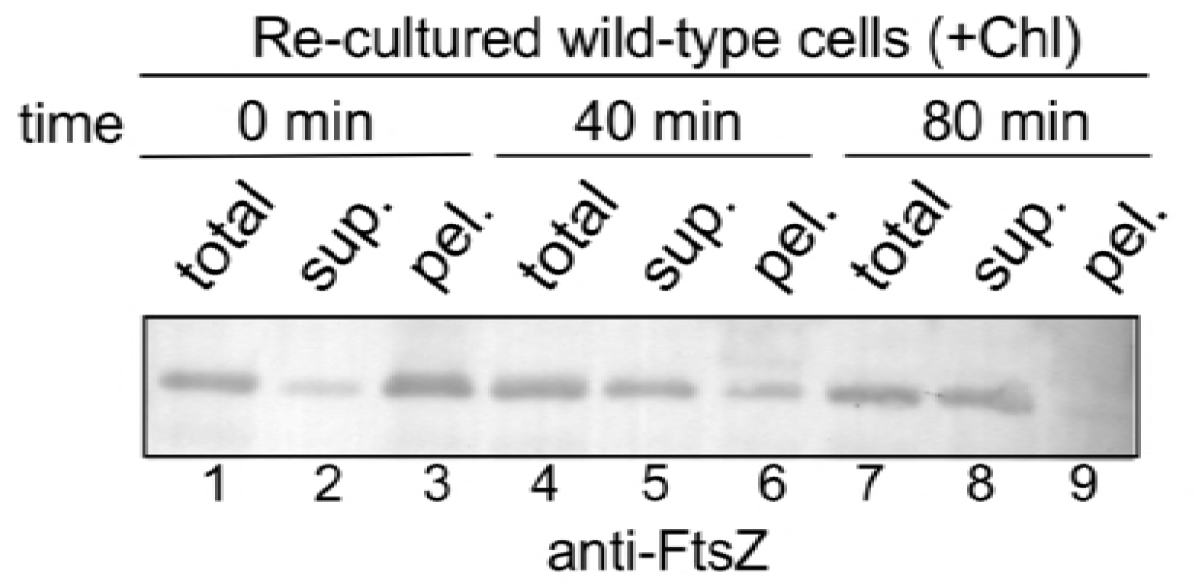
The FtsZ protein disappears from the pellet fraction and reappears in the supernatant fraction in a time-dependent manner when the non-growing late stationary-phase cells are re-cultured in the presence of chloramphenicol. Immunoblotting results for the detection of FtsZ protein in the indicated cell lysate fractions when non-growing late stationary-phase wild-type cells were re-cultured in fresh LB medium containing chloramphenicol to the indicated time points, probed with fresh LB medium containing chloramphenicol to the indicated time points, probed with anti-FtsZ antibodies.

**Figure S7.**
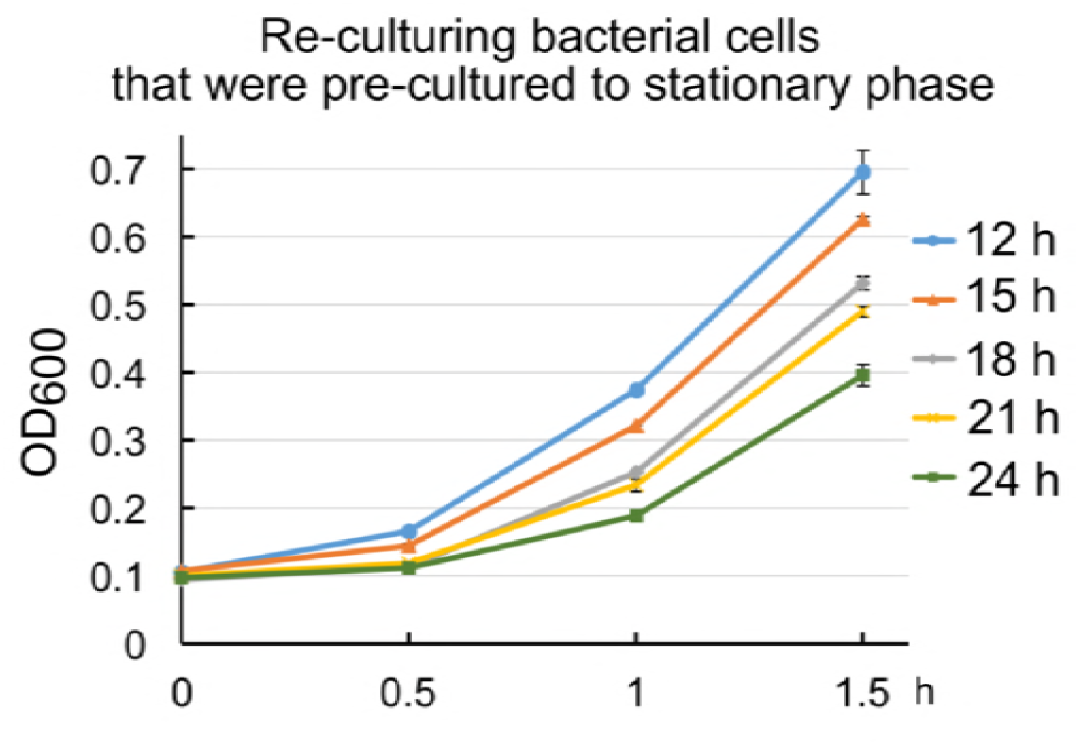
Growth curves of the re-cultured wild-type *E. coli* cells that were precultured to the indicated time points in the stationary phase. These growth curves were used to calculate the average initial doubling time upon redivision (re-division *T*_id_), which reflects the regrowth lag time for each set of the nongrowing stationary phase cells. At least three biological replicates were analyzed for obtaining each value.

**Figure S8.**
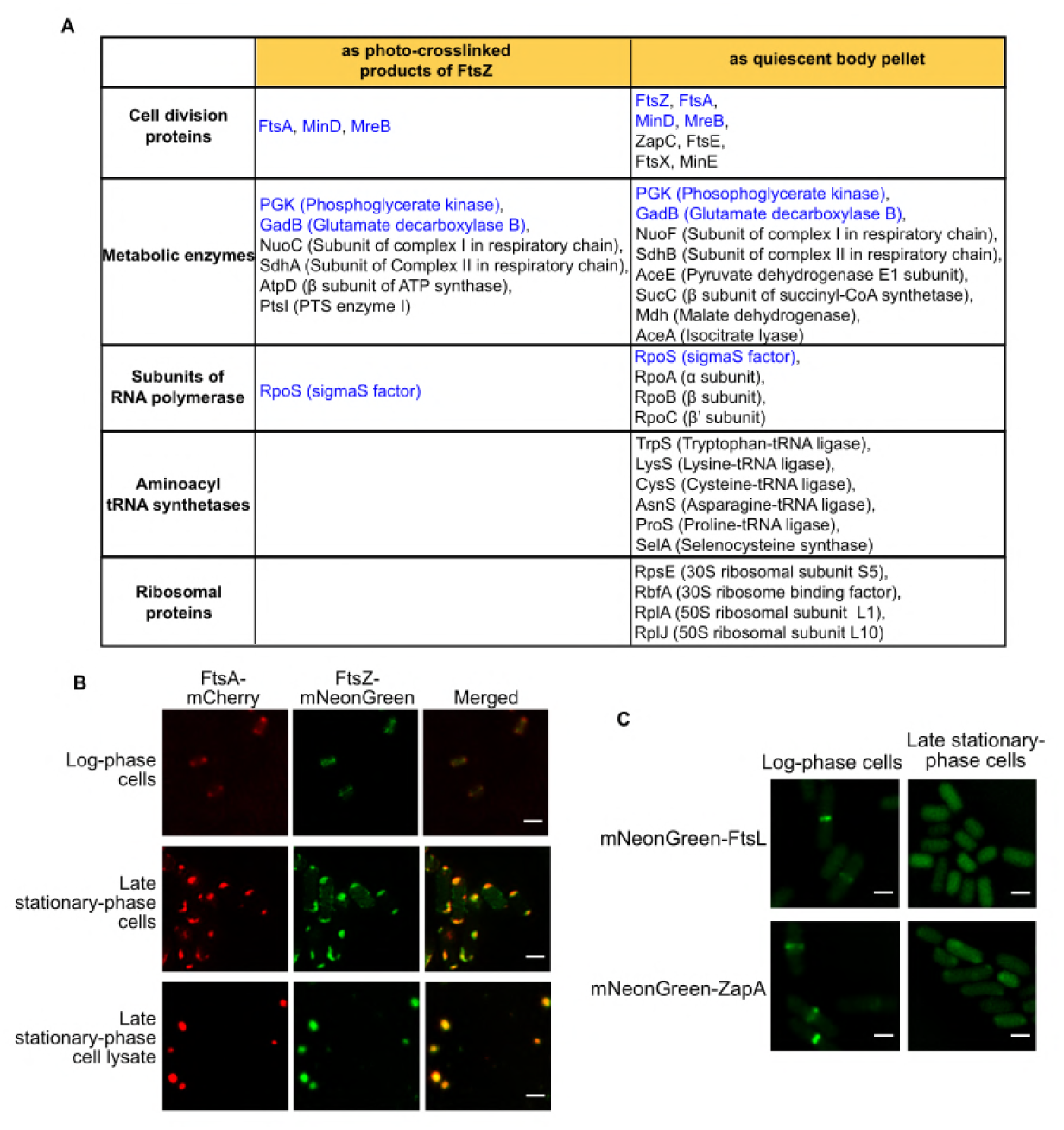
Multiple key cellular proteins are identified in regrowth-delay bodies by mass spectrometry analyses and live-cell imaging analysis verified the presence of FtsA and the absence of FtsL or ZapA in regrowth-delay bodies. **(A)** List of major proteins identified by mass spectrometry analyses, both as the photo-crosslinked products of five pBpa variants of FtsZ (as shown in **Figs. S1A** and **S4D**) and as present in the pellets containing the regrowth-delay bodies, isolated from the non-growing *ftsZ-Avi* and wild-type late stationary-phase cells, respectively. **(B)** Fluorescence microscopic images of the actively dividing log-phase (top) or the non-growing late stationary-phase (middle) *ftsZ-mNeonGreen* cells in which FtsA-mCherry was expressed from a plasmid controlled by a constitutive promoter, as well as of the lysate of the same non-growing late stationary-phase cells (bottom). Scale bars, 1 μm. **(C)** Fluorescence microscopic images of the actively dividing log-phase or nongrowing late stationary-phase cells in which mNeonGreen-FtsL or mNeonGreen-ZapA was heterogeneously expressed from a plasmid. Scale bars, 1 μm.

**Figure S9.**
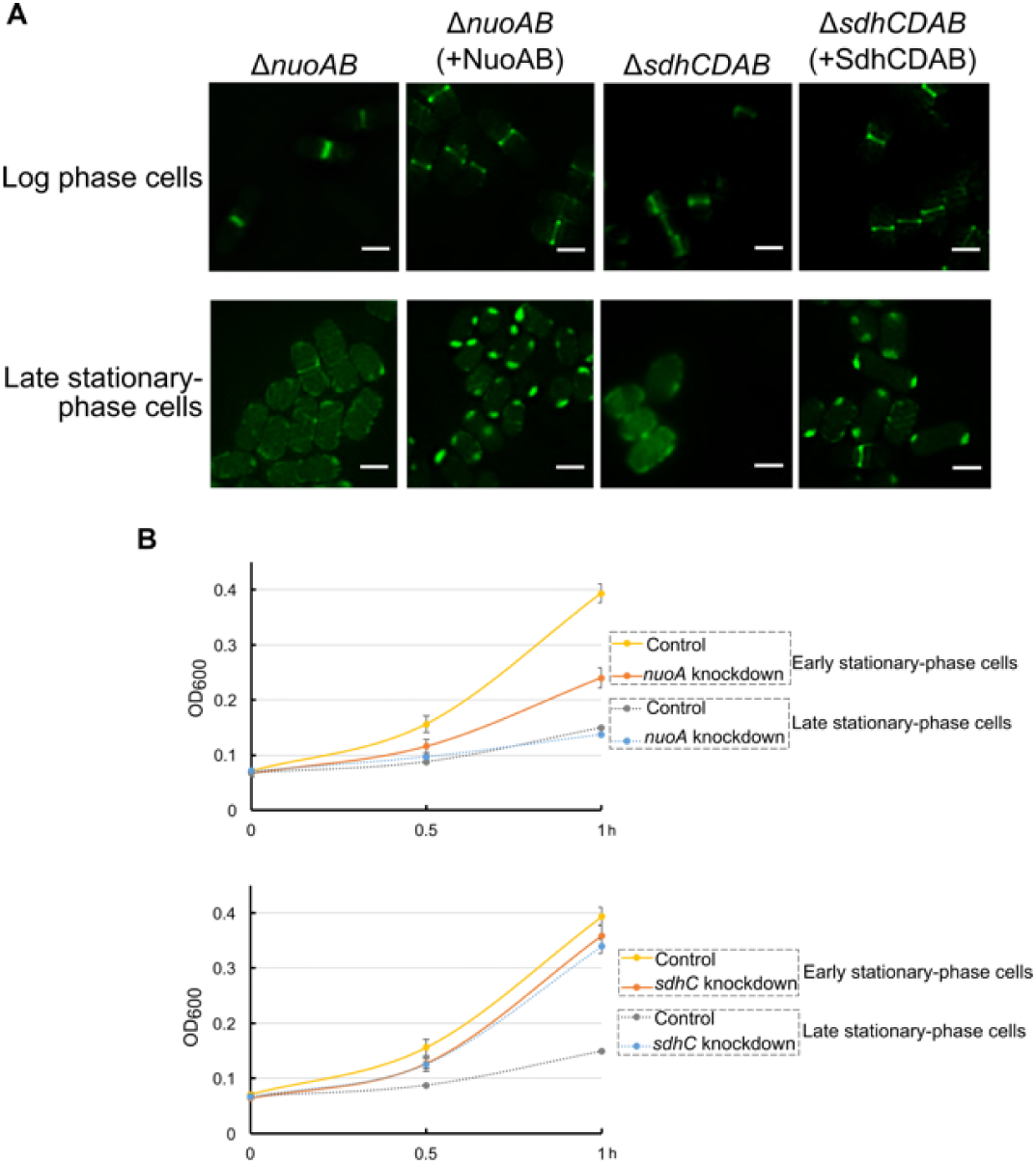
The formation of regrowth-delay bodies in *nuoAB* knockout (*ΔnuoAB*) or *sdhCDAB* knockout (*ΔsdhCDAB*) non-growing late stationary-phase cells is significantly reduced, and the regrowth lag time of non-growing late stationary-phase *ΔnuoA* or *ΔsdhC* knockdown cells is significantly shortened. **(A)** Fluorescence microscopic images of log-phase (top) and late stationary-phase (bottom) *ftsZ-mNeonGreen* cells in which the *nuoAB* or *sdhCDAB* genes were deleted. Scale bars, 1 μm. **(B)** Growth curves of the re-cultured early or late stationary-phase *nuoA* or *sdhC* knockdown cells. Here, cells in which a non-targeting crRNA was expressed from a plasmid were analyzed as the control. All experiments were independently repeated three times.

**Figure S10.**
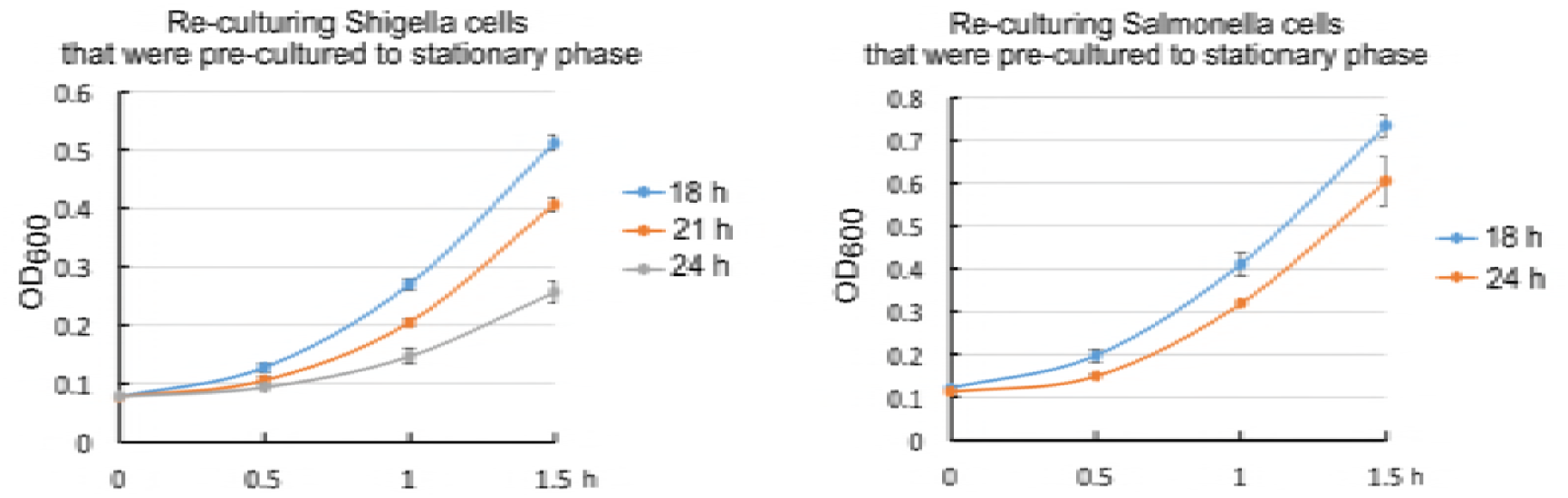
Growth curves of the re-cultured *Shigella* and *Salmonella* bacterial cells that were pre-cultured to the indicated time points in stationary-phase.

## STAR Methods

### Bacterial strains, plasmids, and genome modifications

Listed in **Table S1** are the genotypes of the used *E. coli* strains, all derived from the BW25113 strain with genotype: F-, DE(araD-araB)567, lacZ4787(del)::rrnB-3, LAM-, rph-1, DE(rhaD-rhaB)568, hsdR514 (Blattner et al., 1997). The analyzed pathogenic strains were *Salmonella* Typhimurium SL1344 and *Shigella flexneri* serotype 2a 2457T. All the plasmids employed in this study are listed in **Table S2**. Genome modifications were performed using the λ-red genomic recombination system (Lee et al., 2009). Newly generated plasmids and genome modifications were all confirmed by DNA sequencing.

### Bacterial cell culturing

LB liquid (10 g/l tryptone, 5 g/l yeast extract, and 5 g/l NaCl) and agar-containing solid culture medium were sterilized by autoclaving. *Salmonella* Typhimurium SL1344 and *Shigella flexneri* serotype 2a 2457T were cultured in LB medium with 30 μg/ml streptomycin. For plasmid selection, 50 μg/ml kanamycin, 34 μg/ml chloramphenicol, or 100 μg/ml ampicillin was added to the culture medium. Log-phase and late stationary-phase cells refer to cells that were cultured at 37°C in test tubes and shook at 260 r.p.m. for 6 h and 24 h, respectively, after the overnight-cultured cells were diluted 100-fold in fresh LB medium. The expression of CbtA or KilR was induced by addition of 0.2 μg/ml anhydrotetracycline. For membrane staining, FM4-64 (2 μg/ml) was added to the culture medium, and the cells were then further cultured for another 1 h.

### *In vivo* protein photo-crosslinking of pBpa variants of FtsZ

To perform the photo-crosslinking analysis within the *LY928-ftsZ-Avi* strain (in which endogenous wild-type FtsZ protein was expressed with an Avi tag fused to its C-terminus) that we constructed, each pBpa variant was expressed from a plasmid at a level comparable with that of endogenous FtsZ, and the cells were cultured to log or late stationary phase at 37°C in LB medium containing 200 μM pBpa. The cells were irradiated with UV light (365 nm) for 10 min at room temperature using a Hoefer UVC 500 Crosslinker (Amersham Biosciences, USA) and then collected by centrifugation at 13,000 × *g* before being subjected to further (blotting) analysis.

### Fluorescence microscopic imaging

Cell or cell lysate samples were placed on a glass dish (NEST Biotechnology, USA) and covered with agar before micrographs were acquired at 37°C (for the re-culturing cell samples) or 30°C (for all other samples) with an N-SIM imaging system (Nikon, Japan) using the 2D-SIM mode, a 100×/1.49 NA oil-immersion objective (Nikon, Japan), and under excitation of a 488 nm or 561 nm laser beam. The 3D images were acquired with an N-SIM imaging system using the 3D mode. The samples were sectioned every 120 nm along the Z-axis. The images were further reconstructed using the NIS-Elements AR 4.20.00 (Nikon, Japan) before a further processing with the GNU image manipulation program. At least four images were obtained, and more than 50 bacterial cells were examined for each experiment. All experiments were independently repeated at least three times.

### Cell lysate fractionations

The non-growing late stationary-phase bacterial cells were prepared by growing the cells at 37°C (with shaking at 260 r.p.m.) for 24 h after the overnight-cultured cells were diluted 100-fold into fresh LB medium. The cell samples (such as those used in **Fig. S5**) of the re-culturing experiments were prepared by transferring the 2-fold diluted non-growing late stationary-phase cells into fresh LB medium in the presence of chloramphenicol (34 μg/ml) and further culturing them at 37°C (with shaking at 260 r.p.m.) to the indicated time points. The cells were then collected by centrifugation (8000 × g) and disrupted using a French press at 1000 MPa before centrifugation at 13,000 × *g* to separate the supernatant and pellet fractions.

### Protein purification and mass spectrometry analysis

The photo-crosslinked products of pBpa variants of FtsZ-Avi generated in the LY 928 strain were individually purified using streptavidin magnetic beads after the pellet containing the photo-crosslinked products was dissolved in 8 M urea and diluted 10-fold in binding buffer. The eluted protein samples were then further resolved by SDS-PAGE.

For identification of proteins in the regrowth-delay bodies, the pellet from non-growing late stationary-phase wild type cell lysates was collected, dissolved in 8 M urea, and centrifuged again at 13,000 × *g* before removing the new pellet. The supernatant was then concentrated 10-fold and resolved by SDS-PAGE.

In both of the above cases, the protein bands of SDS-PAGE that could be clearly visualized by Coomassie blue staining on the gel, and were excised and sent for mass spectrometry analysis.

### Blotting analysis

Each sample, including the cell lysate, supernatant fraction, pellet fraction, or UV-irradiated cells, was supplemented with the sample buffer, boiled, and resolved via tricine SDS-PAGE before being further probed with particular antibodies or streptavidin-AP conjugate (for the Avi-tagged proteins) for the blotting analysis. The protein bands visualized on the gels were scanned and processed using the GNU image manipulation program.

### CRISPRi experiments

CRISPRi was performed according to previously reported methods (Luo et al., 2015). Briefly, plasmids carrying a crRNA that targets the *nuoA* or *sdhC* gene were transformed into *E. coli* cells in which the proteins for recognizing and binding specific DNA sequences were expressed from the Cascade operon while the gene *(cas3* gene) encoding the protein that cleaves the target sequence was deleted. The DNA sequences designed for knocking down the *nuoA* and the *sdhC* genes were: AT AGCGAAT GCCCAGT GAT GAGCGAT GACTTC and AATGTGAAAAAACAAAGACCTGTTAATCTGGA, respectively. The control plasmid carried a non-targeting crRNA sequence: CTGCTGGAGCTGGCTG CAAGGCAAGCCGCCCA. The crRNAs on the plasmids were transcribed constitutively rather than induced.

### Cell regrowth and calculation of the average re-division T_id_

Log-phase or late stationary-phase cells of a particular type were diluted 40-fold into fresh LB medium and cultured at 37°C with shaking (260 r.p.m.). Growth curves were prepared by measuring the OD_600_ value of the cultured cells at 30-min intervals. The re-division *T*_id_ value was calculated as 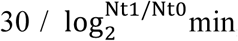, where N_t0_ and N_t1_ were the numbers of cells at 0 min and 30 min, respectively. The N_t1_/N_t0_ ratio for each batch of cultured cells was calculated based on the increase in optical density at 600 nm (the correlation between the cell number and the OD_600_ value was determined by preparing a standard curve). At least three biological replicates were analyzed for obtaining each value.

### Assay for cell survival after antibiotic treatment

Stationary-phase cells were diluted 40-fold into fresh LB medium containing either 5 μg/ml ofloxacin or 200 μg/ml ampicillin and incubated at 37°C with shaking (260 r.p.m.) for 2 h. The cells were then collected by centrifugation (to remove the culture medium and the antibiotics), resuspended in phosphate-buffered saline (PBS), and serially diluted in PBS buffer before being spotted on LB agar plates for CFU counting. The cell survival rate was calculated as follows: [number of colonies formed after antibiotic treatment] / [number of colonies formed without antibiotic treatment] × 100. At least three biological replicates were analyzed for obtaining each value.

**Table S1.** *E. coli* strains used in this study.

| Strain | Genotype<sup>a</sup> | Source/Reference |
| --- | --- | --- |
| BW25113 | $\Delta(araD-araB)567 \Delta lacZ4787(::rrnB-3) rph-1 \Delta(rhaD-rhaB)568 hsdR514$ | (Baba et al., 2006) |
| LY928 | BW25113 $\Delta insH11::aminoacyl-tRNA$ synthetase of pBpa-tRNA<sup>pBpa</sup> | Laboratory storage |
| *ftsZ-Avi* | LY928 *ftsZ::ftsZ-Avi tag* | Laboratory storage |
| *ftsZ-mNeonGreen* | LY928 $\Delta(rhaD-rhaB)568::ftsZ-mNeonGreen$ | Recombineering |
| *ftsZ-mNeonGreen-dnaK-mCherry* | *ftsZ-mNeonGreen dnaK::dnaK-mCherry* | Recombineering |
| *ftsZ-mNeonGreen-clpB-mCherry* | *ftsZ-mNeonGreen clpB::clpB-mCherry* | Recombineering |
| $\Delta cas3$ | LY928 $\Delta cas3 P_{casA}::P_{con}$ | Recombineering (Luo et al., 2015) |
| *ftsZ-mNeonGreen-$\Delta cas3$* | $\Delta cas3 \Delta(rhaD-rhaB)568::ftsZ-mNeonGreen$ | Recombineering |
| $\Delta nuoAB$ | *ftsZ-mNeonGreen $\Delta nuoAB::Kan^R$* | Recombineering |
| $\Delta sdhCDAB$ | *ftsZ-mNeonGreen $\Delta sdhCDAB::Kan^R$* | Recombineering |
| Salmonella | *Salmonella* Typhimurium SL1344 | ATCC |
| Shigella | *Shigella flexneri* serotype 2a 2457T | ATCC |

<sup>a</sup> P<sub>con</sub> is a synthetic constitutive promoter (Luo et al., 2015).

**Table S2.** Plasmids used in this study.

| Plasmid | Genotype<sup>a</sup> | ori | Reference/Source |
| --- | --- | --- | --- |
| pTet-FtsZ-pBpa-mNeonGreen | *bla* P<sub>tet1</sub>::*ftsZ*-*pBpa*-*mNeonGreen* | pBR322 | This study |
| pTet-mNeonGreen | *bla* P<sub>tet1</sub>::*mNeonGreen* | pBR322 | This study |
| pTac-mCherry | *bla* P<sub>con</sub>::*mCherry* | pBR322 | This study |
| pBAD-SSnlpA-mCherry | *bla* P<sub>ara</sub>::*signal peptide of nlpA*-*mCherry* | pBR322 | This study |
| pBAD-OmpA-mCherry | *bla* P<sub>ara</sub>::*ompA*-*mCherry* | pBR322 | This study |
| pACE | *cl* P<sub>ara</sub>::λ-Red recombinase P<sub>ara</sub>::I-SceI endonuclease | p15A | Laboratory storage (Lee et al., 2009) |
| pYLC-rha-FtsZ-mNeonGreen | *bla* upstream homologous sequence-*ftsZ*-*mNeonGreen*-*Kan<sup>R</sup>*- downstream homologous sequence, for inserting *ftsZ*-*mNeonGreen* into the genomic rha operon | pBR322 | This study |
| pYLC-dnaK-mcherry | *bla* upstream homologous sequence-*mCherry*-*Kan<sup>R</sup>*- downstream homologous sequence, for inserting *mCherry* into the C-terminus of genomic *dnaK* | pBR322 | This study |
| pYLC-clpB-mcherry | *bla* upstream homologous sequence-*mCherry*-*Kan<sup>R</sup>*- downstream homologous sequence, for inserting *mCherry* into the C-terminus of genomic *clpB* | pBR322 | This study |
| pTetO-CbtA | *bla* P<sub>tet-T::cbtA-coupling-mCherry-his</sub> | pBR322 | This study |
| pTet-FtsZ-140pBpa | *bla* P<sub>tet1::ftsZ-140pBpa</sub> | pBR322 | Laboratory storage |
| pTet-FtsZ-31pBpa | *bla* P<sub>tet1::ftsZ-31pBpa</sub> | pBR322 | Laboratory storage |
| pTet-FtsZ-47pBpa | *bla* P<sub>tet1::ftsZ-47pBpa</sub> | pBR322 | Laboratory storage |
| pTet-FtsZ-51pBpa | *bla* P<sub>tet1::ftsZ-51pBpa</sub> | pBR322 | Laboratory storage |
| pTet-FtsZ-54pBpa | *bla* P<sub>tet1::ftsZ-54pBpa</sub> | pBR322 | Laboratory storage |
| pTet-FtsZ-61pBpa | *bla* P<sub>tet1::ftsZ-61pBpa</sub> | pBR322 | Laboratory storage |
| pTet-FtsZ-82pBpa | *bla* P<sub>tet1::ftsZ-82pBpa</sub> | pBR322 | Laboratory storage |
| pTet-FtsZ-85pBpa | *bla* P<sub>tet1::ftsZ-85pBpa</sub> | pBR322 | Laboratory storage |
| pTet-FtsZ-114pBpa | *bla* P<sub>tet1::ftsZ-114pBpa</sub> | pBR322 | Laboratory storage |
| pTet-FtsZ-151pBpa | *bla* P<sub>tet1::ftsZ-151pBpa</sub> | pBR322 | Laboratory storage |
| pTet-FtsZ-166pBpa | *bla* P<sub>tet1::ftsZ-166pBpa</sub> | pBR322 | Laboratory storage |
| pTet-FtsZ-174pBpa | *bla* P<sub>tet1::ftsZ-174pBpa</sub> | pBR322 | Laboratory storage |
| pTet-FtsZ-288pBpa | *bla* P<sub>tet1::ftsZ-288pBpa</sub> | pBR322 | Laboratory storage |
| pTet-FtsZ-340pBpa | *bla* P<sub>tet1::ftsZ-340pBpa</sub> | pBR322 | Laboratory storage |
| pTet-FtsZ-348pBpa | *bla* P<sub>tet1::ftsZ-348pBpa</sub> | pBR322 | Laboratory storage |
| pTac-trpS-Avi | *bla* P<sub>con::trpS-Avitag</sub> | pBR322 | This study |
| pTac-rpoS-Avi | *bla* P<sub>con::rpoS-Avitag</sub> | pBR322 | This study |
| pTac-aceE-Avi | *bla* P<sub>con::aceE-Avitag</sub> | pBR322 | This study |
| pTac-rplL-Avi | *bla* P<sub>con::rplL-Avitag</sub> | pBR322 | This study |
| pTac-FtsA-mNeonGreen | *bla* P<sub>con</sub>::*ftsA-mNeonGreen*<br>(Durand-Heredia et al., 2011) | pBR322 | This study |
| pTac-FtsA-mCherry | *bla* P<sub>con</sub>::*ftsA-mCherry*<br>(Ma et al., 1996) | pBR322 | This study |
| pTac-mNeonGreen-ZapA | *bla* P<sub>con</sub>::*zapA-mNeonGreen* | pBR322 | This study |
| pTac-ZapC-mNeonGreen | *bla* P<sub>con</sub>::*zapC-mNeonGreen* | pBR322 | This study |
| pTac-mNeonGreen-FtsL | *bla* P<sub>con</sub>:: *mNeonGreen-ftsL*<br>(Ghigo and Beckwith, 2000) | pBR322 | This study |
| pYLC-Δ*Cas3*-kana | *bla* upstream homologous<br>sequence-*Kan<sup>R</sup>*-P<sub>con</sub>-downstream<br>homologous sequence, for<br>deleting the *cas3* gene and<br>replacing the native promoter of<br>the Cascade operon with a<br>constitutive promoter | pBR322 | This study |
| pTac-nuoAi | *cat* P<sub>con</sub>::crRNA sequence<br>targeting *nuoA* | pBR322 | This study |
| pTac-sdhCi | *cat* P<sub>con</sub>::crRNA sequence<br>targeting *sdhC* | pBR322 | This study |
| pYLC-Δ*nuoAB*-kana | *bla* upstream homologous<br>sequence-*Kan<sup>R</sup>*- downstream<br>homologous sequence, for<br>deleting the *nuoAB* gene | pBR322 | This study |
| pYLC-Δ*sdhCDAB*-kana | *bla* upstream homologous<br>sequence-*Kan<sup>R</sup>*- downstream<br>homologous sequence, for<br>deleting the *sdhCDAB* gene | pBR322 | This study |
| pTac-NuoAB | *bla* P<sub>con</sub>::*nuoAB* | pBR322 | This study |
| pTac-sdhCDAB | *bla* P<sub>con</sub>::*sdhCDAB* | pBR322 | This study |

<sup>a</sup> P<sub>tet</sub>, P<sub>ara</sub> and P<sub>con</sub> indicate the Tet-on/Tet-off, arabinose and synthetic constitutive (selected from the Anderson promoter collection: parts.igem.org/Promoters/Catalog/Anderson) promoters, respectively. P<sub>tet1</sub> indicates that the expression of proteins was not induced by anhydrotetracycline, just via leaky expression. P<sub>tet-T</sub> indicates that the λt1 transcriptional terminator was inserted before the Tet promoter to achieve a stringent expression.

